# Origins and Evolution of the Global RNA Virome

**DOI:** 10.1101/451740

**Authors:** Yuri I. Wolf, Darius Kazlauskas, Jaime Iranzo, Adriana LucÍa-Sanz, Jens H. Kuhn, Mart Krupovic, Valerian V. Dolja, Eugene V. Koonin

## Abstract

Viruses with RNA genomes dominate the eukaryotic virome, reaching enormous diversity in animals and plants. The recent advances of metaviromics prompted us to perform a detailed phylogenomic reconstruction of the evolution of the dramatically expanded global RNA virome. The only universal gene among RNA viruses is the RNA-dependent RNA polymerase (RdRp). We developed an iterative computational procedure that alternates the RdRp phylogenetic tree construction with refinement of the underlying multiple sequence alignments. The resulting tree encompasses 4,617 RNA virus RdRps and consists of 5 major branches, 2 of which include positive-sense RNA viruses, 1 is a mix of positive-sense (+) RNA and double-stranded (ds) RNA viruses, and 2 consist of dsRNA and negative-sense (−) RNA viruses, respectively. This tree topology implies that dsRNA viruses evolved from +RNA viruses on at least two independent occasions, whereas -RNA viruses evolved from dsRNA viruses. Reconstruction of RNA virus evolution using the RdRp tree as the scaffold suggests that the last common ancestors of the major branches of +RNA viruses encoded only the RdRp and a single jelly-roll capsid protein. Subsequent evolution involved independent capture of additional genes, particularly, those encoding distinct RNA helicases, enabling replication of larger RNA genomes and facilitating virus genome expression and virus-host interactions. Phylogenomic analysis reveals extensive gene module exchange among diverse viruses and horizontal virus transfer between distantly related hosts. Although the network of evolutionary relationships within the RNA virome is bound to further expand, the present results call for a thorough reevaluation of the RNA virus taxonomy.

**IMPORTANCE:** The majority of the diverse viruses infecting eukaryotes have RNA genomes, including numerous human, animal, and plant pathogens. Recent advances of metagenomics have led to the discovery of many new groups of RNA viruses in a wide range of hosts. These findings enable a far more complete reconstruction of the evolution of RNA viruses than what was attainable previously. This reconstruction reveals the relationships between different Baltimore Classes of viruses and indicates extensive transfer of viruses between distantly related hosts, such as plants and animals. These results call for a major revision of the existing taxonomy of RNA viruses.

## Introduction

Early evolution of life is widely believed to have first involved RNA molecules that functioned both as information storage devices and as catalysts (ribozymes) (1, 2). Subsequent evolution involved the emergence of DNA, the dedicated genomic material, and proteins, the ultimate operational molecules. RNA molecules remained central for translating information from genes to proteins (mRNA), the functioning of the translation machinery itself (rRNA and tRNA), and for a variety of regulatory functions (various classes of non-coding RNA that are increasingly discovered in all life forms) (3). Viruses with RNA genomes (“RNA viruses” for short) that do not involve DNA in their genome replication and expression cycles (4, 5) can be considered the closest extant recapitulation, and possibly, a relic of the primordial RNA world.

RNA viruses comprise 3 of the 7 so-called Baltimore Classes of viruses that differ with respect to the nature of the genome (i.e., the nucleic acid form that is packaged into virions) and correspond to distinct strategies of genome replication and expression: positive-sense (+) RNA viruses, double-stranded (ds) RNA viruses and negative-sense (−) RNA viruses (6). The +RNA viruses use the simplest possible strategy of replication and expression as the same molecule functions as both genome and mRNA (7). Most likely, the first replicators to emerge in the RNA world, after the evolution of translation, resembled +RNA viruses (8). Because the +RNA released from a virion can be directly used for translation to produce viral proteins, virions of +RNA viruses contain only structural proteins, in addition to the genome. In contrast, -RNA and dsRNA viruses package their transcription and replication machineries into their virions because these are necessary to initiate the virus reproduction cycles (9, 10).

RNA viruses comprise a major part of the global virome. In prokaryotes, the known representation of RNA viruses is narrow. Only one family of +RNA viruses *(Leviviridae)* and one family of dsRNA viruses *(Cystoviridae)* are formally recognized, and furthermore, their members have limited host ranges. No-RNA viruses have been isolated from prokaryotes (4, 11, 12). Although recent metagenomic studies suggest that genetic diversity and host range of prokaryotic RNA viruses could be substantially underestimated (13, 14), it appears that the scope of the prokaryotic RNA virome is incomparably less than that of the DNA virome. As discussed previously, potential causes of the vast expansion of the RNA virome in eukaryotes might include the emergence of the compartmentalized cytosol that provided a hospitable, protective environment for RNA replication that is known to be associated with the endoplasmic reticulum and other membrane compartments (11). Conversely, the nuclear envelope could be a barrier that prevents the access of DNA viruses to the host replication and transcription machineries and thus partially relieves the stiff competition that DNA viruses of prokaryotes represent for RNA viruses.

In a sharp contrast, the 3 Baltimore Classes of RNA viruses dominate the eukaryotic virosphere (11, 15). Eukaryotes from all major taxa are hosts to RNA viruses, and particularly in plants and invertebrates, these viruses are enormously abundant and diverse (15-17). Until recently, the study of RNA viromes had been heavily skewed towards viruses infecting humans, livestock, and agricultural plants. Because of these limitations and biases, the attempts to reconstruct the evolutionary history of RNA viruses were bound to be incomplete. Nonetheless, these studies have yielded important generalizations. A single gene encoding an RNA-dependent RNA polymerase (RdRp) is universal among RNA viruses, including capsid-less RNA replicons but excluding some satellite viruses (18). Even within each of the 3 Baltimore Classes, virus genomes do not include fully conserved genes other than those encoding RdRps (15). However, several hallmark genes are shared by broad ranges of RNA viruses including, most notably, those encoding capsid proteins of icosahedral and helical virions of +RNA viruses (19, 20), and key enzymes involved in virus replication such as distinct helicases and capping enzymes (15).

The RdRp gene and encoded protein are natural targets of evolutionary analysis because they are the only universal gene (protein) in RNA viruses. However, obtaining strongly supported phylogenies for RdRps is a difficult task due to the extensive sequence divergence, apart from several conserved motifs that are required for polymerase activity (21-23). RdRps belong to the expansive class of polymerases containing so-called Palm catalytic domains along with the accessory Fingers and Thumb domains (24, 25). In addition to viral RdRps, Palm domain polymerases include reverse transcriptases (RTs) of retroelements and reverse-transcribing viruses, and DNA polymerases that are responsible for genome replication in cellular organisms and diverse DNA viruses. Within the Palm domain class of proteins, RT and the +RNA virus RdRps are significantly similar in sequence and structure, and appear to comprise a monophyletic group (22, 24-26). More specifically, the highest similarity is observed between the +RNA virus RdRps and the RTs of group II introns. These introns are wide-spread retrotransposons in prokaryotes that are thought to be ancestral to the RTs of all other retrotransposons as well as retroviruses and pararetroviruses (recently jointly classified as the order *Ortervirales)* of eukaryotes (27-30).

A phylogenetic analysis of the +RNA virus RdRps revealed only a distant relationship between the leviviruses and the bulk of the eukaryotic +RNA viruses, leaving the ancestral relationships uncertain (31). The origin of eukaryotic +RNA viruses from their prokaryotic counterparts is an obvious possibility. However, given the dramatically greater prevalence of +RNA viruses among eukaryotes compared to the narrow spread of leviviruses and “levi-like viruses” in bacteria, an alternative scenario has been proposed. Under this scenario, RdRps of the prokaryotic and eukaryotic +RNA viruses independently descended from distinct RTs (11, 31). Among eukaryotic +RNA viruses, phylogenetic analysis of the RdRps strongly supports the existence of picornavirus and alphavirus “supergroups”, which are further validated by additional signature genes (7, 15). However, both the exact compositions of these supergroups and the evolutionary relationships among many additional groups of viruses remain uncertain. Some RdRp phylogenies suggest a third supergroup combining animal “flavi-like viruses” and plant tombusviruses but this unification is not supported by additional shared genes and thus remains tenuous (7, 15, 21).

The similarity of RdRps among dsRNA viruses is limited but these RdRps are similar to varying degree to +RNA virus RdRps. Therefore, different groups of dsRNA viruses might have evolved from different +RNA viruses independently, on multiple occasions (15, 32). Although it is not entirely clear how prokaryotic dsRNA viruses fit into this concept, evidence of an evolutionary affinity between cystoviruses and reoviruses has been presented (33, 34).

For a long time, the evolutionary provenance of-RNA virus RdRps remained uncertain due to their low sequence similarity to other RdRps and RTs (23). However, recent protein structure comparisons point to a striking similarity between the RdRps of-RNA orthomyxoviruses and those of +RNA flaviviruses and dsRNA cystoviruses (35). All these findings notwithstanding, the overall evolutionary relationships among the RdRps of +RNA,-RNA, and dsRNA viruses, and RTs remain unresolved. In particular, whether the RdRps of +RNA and-RNA viruses are mono-or polyphyletic, is unclear.

Many deep evolutionary connections between RNA virus groups that originally were thought to be unrelated have been delineated using the results of pre-metagenomic era evolutionary studies. These discoveries culminated in the establishment of RNA virus supergroups (7, 9, 36).

However, the evolutionary provenance of many other RNA virus groups remained unclear, and so did the relationships between the RNA viruses of the 3 Baltimore Classes and retroelements, and their ultimate origins. The prospects of substantial progress appeared dim because of the extreme sequence divergence among RNA viruses, which could amount to irrevocable loss of evolutionary information.

Recent revolutionary developments in virus metagenomics (metaviromics) dramatically expanded the known diversity of RNA viruses and provided an unprecedented amount of sequence data for informed investigation into RNA virus evolution (11, 17, 37). The foremost development was the massive expansion of the known invertebrate virome that was achieved primarily through meta-transcriptome sequencing of various holobionts. The subsequent phylogenetic analysis revealed previously unknown lineages of +RNA and-RNA viruses and prompted reconsideration of high rank virus unifications, such as +RNA virus supergroups (14, 38-41). The RNA viromes of fungi and prokaryotes also underwent notable expansion albeit not as massive as that of invertebrates (13, 42, 43).

Here we reexamine the evolutionary relationships among and within the 3 Baltimore Classes of RNA viruses through a comprehensive analysis of the available genomic and metagenomic sequences. In particular, to build a phylogenetic tree of thousands of viral RdRps, we designed an iterative computational procedure that alternates phylogeny construction with refinement of the underlying multiple alignments. Although RNA viruses have relatively short genomes (~3-41 kb), the combined gene repertoire (pangenome) of these viruses includes numerous genes that are shared, to varying degrees, by related subsets of RNA viruses. To obtain further insight into virus evolution, we therefore attempted to reconstruct the history of gain and loss of conserved proteins and domains in different virus lineages. We also investigated evolution of the single jelly-roll capsid protein (SJR-CP), the dominant type of capsid protein among +RNA viruses. Our analysis revealed patterns that are generally congruent with the RdRp phylogeny and provide further insights into the evolution of different branches of RNA viruses. Finally, we analyzed a bipartite network in which RNA virus genomes are connected via nodes representing virus genes (44, 45) to identify distinct modules in the RNA virosphere. The results shed light on the evolution of RNA viruses, revealing, in particular, the monophyly of-RNA viruses and their apparent origin from dsRNA viruses, which seem to have evolved from distinct branches of +RNA viruses on at least two independent occasions.

## RESULTS

### Comprehensive phylogeny of RNA virus RNA-dependent RNA polymerases: overall structure of the tree and the 5 major branches

Amino acid sequences of RdRps and RTs were collected from the non-redundant National Center for Biotechnology Information (NCBI) database and analyzed using an iterative clustering-alignment-phylogeny procedure (Figure S1; see Materials and Methods for details). This procedure ultimately yielded a single multiple alignment of the complete set of 4,617 virus RdRp sequences (Data set S1) and 1,028 RT sequences organized in 50+2 clusters (50 clusters of RdRps and 2 clusters of RTs; see Materials and Methods for details). This sequence set did not include RdRps of members of the families *Birnaviridae* or *Permutatetraviridae,* distinct groups of RNA viruses that encompass a circular permutation within the RdRp Palm domain (46) and therefore could not be confidently included in the alignment over their entire lengths.

The phylogenetic tree of RdRps and RTs (Data set S2 and Figure 1) was assembled from a set of trees that represent three hierarchical levels of relationships. At the lowest level, full complements of sequences from each cluster were used to construct cluster-specific trees. At the intermediate level, up to 15 representatives from each cluster were selected to elucidate supergroup-level phylogeny. At the highest level, up to 5 representatives from each cluster were taken to resolve global relationships (Data set S3A and S4). The final tree (Data set S2) was assembled by replacing the cluster representatives with the trees from the previous steps.

**Figure.1.**
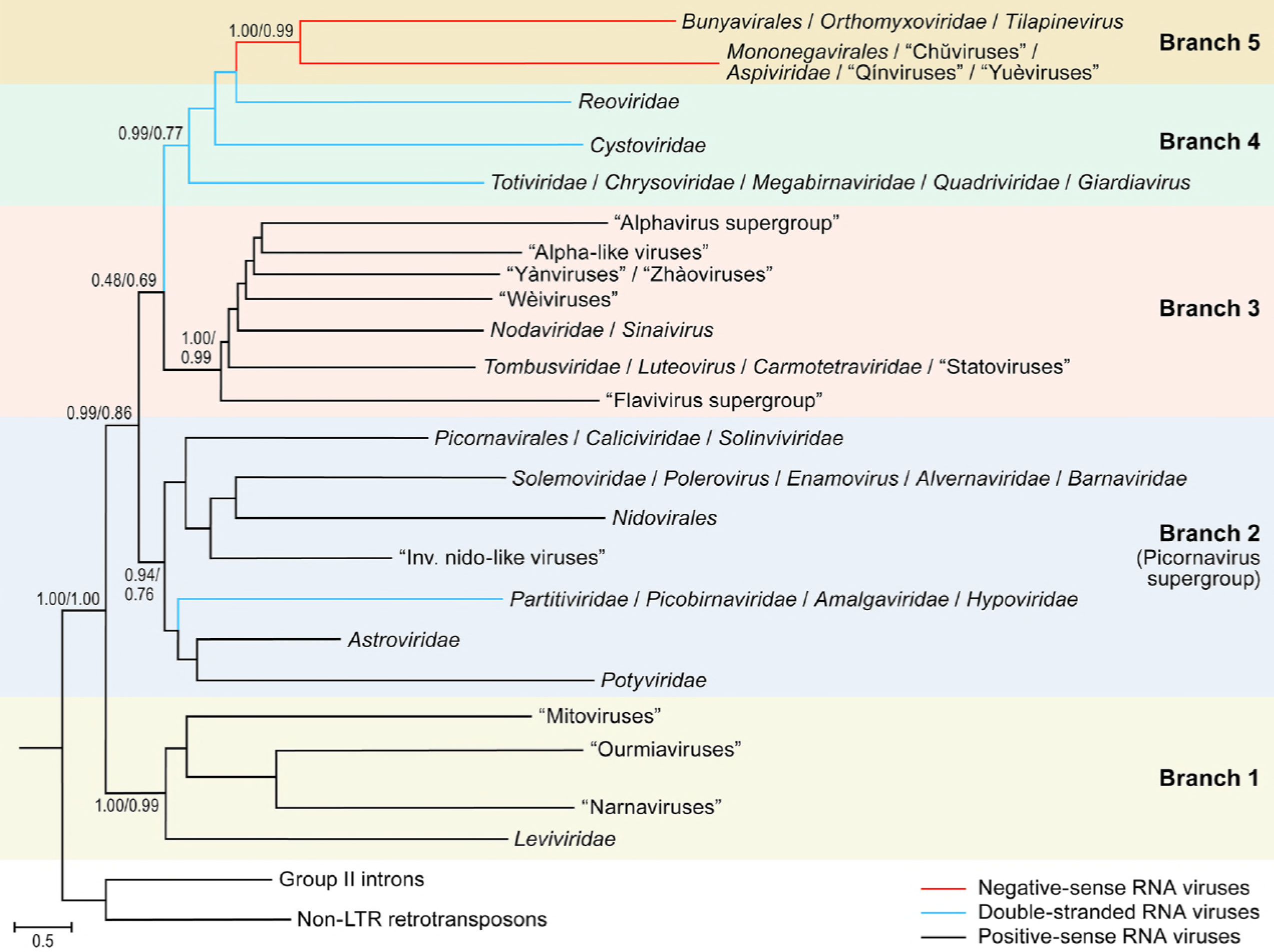
Phylogeny of RNA virus RNA-dependent RNA polymerases (RdRps) and reverse transcriptases (RTs): the main branches (1-5). Each branch represents collapsed sequences of the respective set of RdRps. The 5 main branches discussed in the text are labeled accordingly. The bootstrap support obtained by (numerator) and by (denominator) is shown for each internal branch. LTR, long-terminal repeat.

The large number and immense diversity of the viruses included in our analysis create serious challenges for a systematic, phylogeny-based nomenclature of the identified evolutionary lineages of RNA viruses. Many such lineages consist of viruses newly discovered by metaviromics, are not yet formally classified by the International Committee on Taxonomy of Viruses (ICTV), and therefore, cannot be assigned formal names. For the purpose of the present work, we adopted a semi-arbitrary naming scheme using the following approach. 1) We use taxon names that have been fully accepted by the ICTV as of March 2018 (47) whenever possible. These names are recognizable through their capitalization and italicization and rank-specific suffixes (e.g., *-virales* for orders, *-viridae* for families). As is the common practice in virus taxonomy, the officially classified members of each ICTV-approved taxon are referred to via vernaculars (recognizable through their lack of capitalization and italicization). For instance, the members of the ICTV-approved order *Bunyavirales* are called bunyaviruses, whereas those of the family *Tombusviridae* are called tombusviruses. However, in this work, both taxon and vernacular terms are to be understood *sensu lato:* if our analysis indicates certain viruses to be members of or very closely related to an ICTV-established taxon, we consider them members of that taxon despite the lack of current ICTV recognition. As a result, in our analysis, the order *Bunyavirales* has more members than in the official taxonomy. 2) we use vernacular names in quotation marks for viruses/lineages that are clearly distinct from those covered by the official ICTV framework. Whenever possible, we use names that circulate in the literature (e.g., “hepeliviruses”, “statoviruses”). In the absence of such unofficial names, we name the lineage reminiscent of the next closely related lineage (e.g., “levi-like viruses” are a clearly distinct sister group to *Leviviridae*/leviviruses). 3) Monophyletic clusters that transcend the currently highest ICTV-accepted rank (i.e., order) are labeled according to terms circulating in the literature (i.e., “alphavirus supergroup”, “flavivirus supergroup”, and “picornavirus supergroup”). 4) Lineages represented by a single virus are labeled with the respective virus name.

Rooting the phylogenetic tree, generated using PhyML (48), between the RTs and RdRps resulted in a well-resolved topology of RNA viruses which the tree splits into 5 major branches, each including substantial diversity of viruses (Figure 1).

Branch 1: leviviruses and their eukaryotic relatives, namely, “mitoviruses”, “narnaviruses” and “ourmiaviruses” (the latter three terms are placed in quotation marks as our analysis contradicts the current ICTV framework, which considers mitoviruses and narnaviruses members of one family, *Narnaviridae,* and ourmiaviruses members of a free-floating genus *Ourmiavirus);*

Branch 2 (“picornavirus supergroup”): a large assemblage of +RNA viruses of eukaryotes, in particular, those of the orders *Picornavirales* and *Nidovirales,* the families *Caliciviridae, Potyviridae, Astroviridae,* and *Solemoviridae,* a lineage of dsRNA viruses including partitiviruses and picobirnaviruses; and several other, smaller groups of +RNA and dsRNA viruses.

Branch 3: a distinct subset of +RNA viruses including the “alphavirus supergroup” along with “flavivirus supergroup”, nodaviruses, and tombusviruses, the “statovirus”, “weivirus”, “yanvirus”, “zhàovirus” groups; and several additional, smaller groups.

Branch 4: dsRNA viruses including cystoviruses, reoviruses, and totivirues, and several additional families.

Branch 5:-RNA viruses.

Each of these 5 major branches of the tree is strongly supported by bootstrap replications (Figure 1). Assuming the RT rooting of the tree, Branch 1, which consists of leviviruses and their relatives infecting eukaryotes, is a sister group to the rest of RNA viruses; this position is highly robust to the choice of the phylogenetic method and parameters. This tree topology is compatible with the monophyly of the RdRps and, by inference, of RNA viruses and the origin of eukaryotic RNA viruses from a prokaryotic RNA virus ancestor shared with leviviruses. The deeper history remains murky. We have no information on the nature of the common ancestor of retroelements and RNA viruses, let alone whether the ancestor was an RNA virus or a retroelement. However, parsimony considerations suggest that a retroelement ancestor is more likely given that capsids first appeared in the virus part of the tree rather than having been lost in retroelements.

The next split in the tree occurs between Branch 2 and the short stem that formally joins Branches 3, 4, and 5. However, the unification of Branch 3 with Branches 4 and 5 is weakly supported and might not reflect actual common ancestry.

Arguably, the most striking feature of the RNA virus tree topology is the paraphyly of +RNA viruses relative to dsRNA and-RNA viruses. Indeed, according to this phylogeny,-RNA viruses evolved from within dsRNA viruses, whereas dsRNA viruses are polyphyletic (Figure 1). One major group of dsRNA viruses that includes partitiviruses and picobirnaviruses is firmly embedded within the +RNA virus Branch 2, whereas another, larger dsRNA virus group including cystoviruses, reoviruses, totiviruses and viruses from several other families comprise the distinct Branch 4 that might be related to +RNA virus Branch 3 (Figure 1). This placement of the two branches of dsRNA viruses is conceptually compatible with the previous evolutionary scenarios of independent origins from +RNA viruses. However, the presence of a strongly supported branch combining 3 lineages of dsRNA viruses that infect both prokaryotes and eukaryotes suggests a lesser extent of polyphyly in the evolution of dsRNA viruses than originally proposed (15, 32).

An alternative phylogenetic analysis of the same RdRp alignment using RAxML yielded the same 5 main branches, albeit some with weak support (Data set S3B). Furthermore, although the dsRNA viruses are split the same way using RAxML as in the PhyML tree, the nested tree topology, in which Branch 4 (the bulk of dsRNA viruses) is lodged deep within +RNA viruses, and Branch 5 (-RNA viruses) is located inside Branch 4, is not reproduced (Data set S3B). Instead, Branches 4 and 5 are separate and positioned deep in the tree, right above the split between Branch 1 and the rest of the RdRps. Given the poor resolution of the RAxML tree and a strong biological argument, namely, the absence of identified-RNA viruses in prokaryotes or protists (with the exception of the “leishbuviruses” infecting kinetoplastids (49, 50); see also discussion below), we believe that the tree topology in Figure 1 carries more credence than that shown in Data set S3B. Nevertheless, these discrepancies emphasize that utmost caution is due when biological interpretation of deep branching in trees of highly divergent proteins is attempted.

### Evolution of the 5 major branches of RNA viruses and reconstruction of gene gain and loss events

#### Reconstruction of gene gains and losses

The RdRp is the only universal protein of RNA viruses. Accordingly, other viral genes must have been gained and/or lost at different stages of evolution. Thus, after performing the phylogenetic analysis of RdRps, we assigned the proteins and domains shared by diverse viruses to the branches of the RdRp tree. Multiple alignments and hidden Markov model (hmm) profiles were constructed for 16,814 proteins and domains encoded by RNA viruses, and a computational pipeline was developed to map these domains on the viral genomes (Data set S5). The resulting patterns of domain presence/absence in the branches of the RdRp tree were used to reconstruct the history of the gains and losses of RNA virus genes (or proteins and domains, for simplicity), both formally, using the ML-based Gloome method (51), and informally, from parsimony considerations.

These reconstructions reveal a high level of branch-specificity in the evolution of the gene repertoire of RNA viruses. The only protein that is likely to have been gained at the base of the eukaryotic virus subtree (Branches 2 to 5) (that also includes the bacterial cystoviruses) is the single jelly-roll capsid protein (SJR-CP) (Figure S2A). Retroelements lack capsid proteins, and therefore, there is no indication that SJR-CP was present in the hypothetical element that encoded the common ancestor of RdRps and RTs. Furthermore, reconstruction of the evolution of Branch 1 (leviviruses and their relatives) argues against the ancestral status of SJR-CP in this branch.

#### Branch 1: leviviruses and their descendants

The current RdRp tree topology combined with gene gain-loss reconstruction suggests the following evolutionary scenario for Branch 1 (Figure 2A): a levivirus-like ancestor that, like the extant members of the *Leviviridae,* possessed a capsid protein unrelated to SJR-CP (19, 52) gave rise to naked eukaryotic RNA replicons known as “mitoviruses” and “narnaviruses”. These replicons consist of a single RdRp gene (Fig. 2B) and replicate either in mitochondria or in the cytosol of the host cells, respectively, of fungal and invertebrate hosts (the latter hosts were identified in metaviromic holobiont analyses) (14, 53). Recently, the existence of plant “mitoviruses” has been reported although it is not known whether these viruses reproduce in the mitochondria (54). The “narnavirus” RdRp is also the ancestor of the RdRp of the expanding group of “ourmiaviruses” (Figure 2A). “Ourmiaviruses” were originally identified in a narrow range of plants, and genomic analysis revealed the chimeric nature of these viruses, with a “narnavirus”-like RdRp but SJR-CPs and movement proteins (MPs) which were apparently acquired from “picorna-like” and “tombus-like” viruses, respectively (55). Use of metaviromics has led to the identification of numerous related viruses associated with invertebrates, many of which encode distinct SJR-CP variants and some of which acquired an RNA helicase (Figure 2B and S2) (14). Thus, the evolution of this branch apparently involved the loss of the structural module of leviviruses, which yielded naked RNA replicons that reproduced in the mitochondria of early eukaryotes. A group of these replicons subsequently escaped to the cytosol which was followed by the reacquisition of unrelated structural modules from distinct lineages of eukaryotic viruses inhabiting the same environment (Figure 2B). This complex evolutionary scenario emphasizes the key role of modular gene exchange in the evolution of RNA viruses.

**Figure.2.**
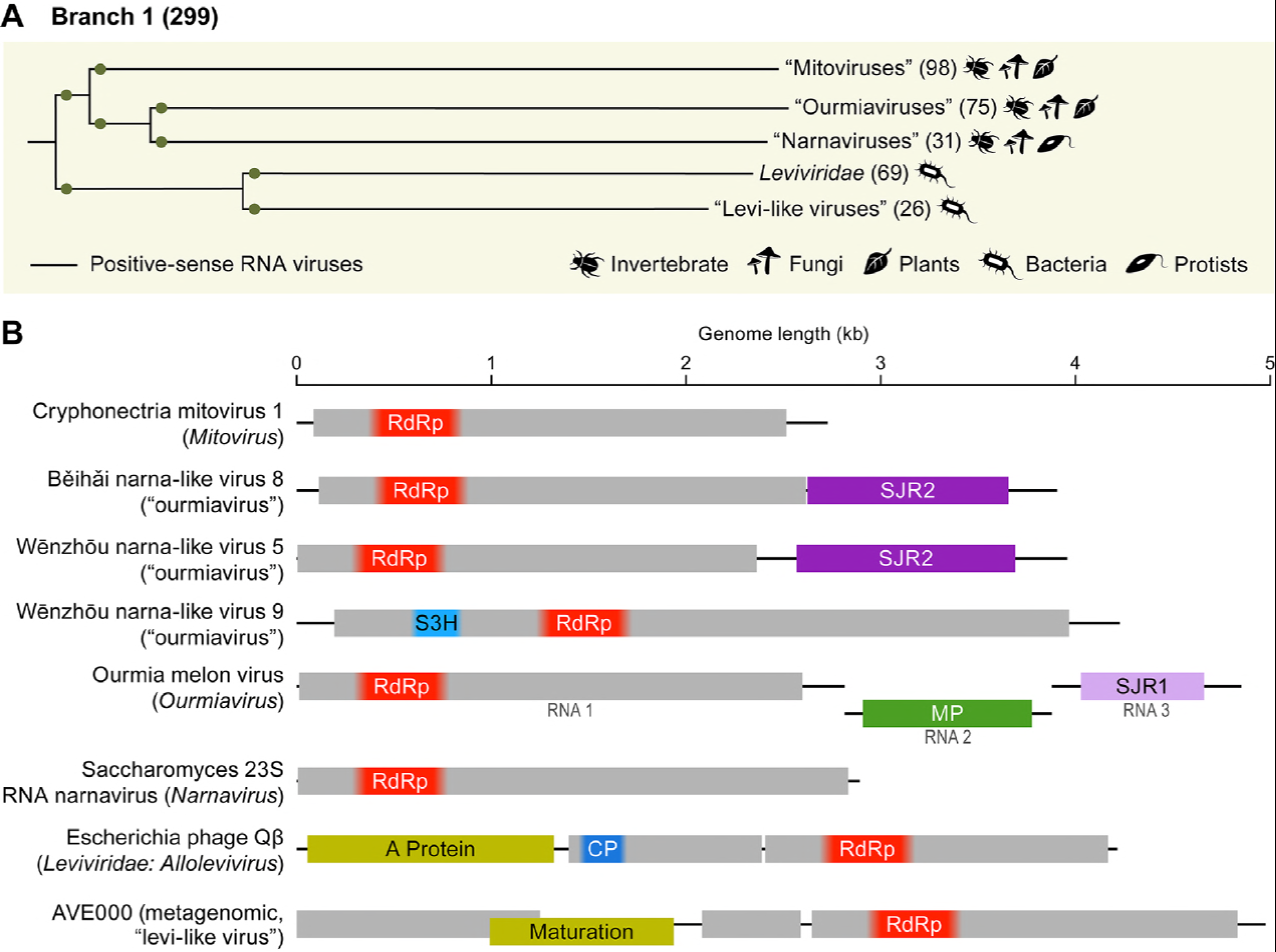
Branch 1 of the RNA virus RNA-dependent RNA polymerases (RdRps): leviviruses and their relatives. (A) Phylogenetic tree of the virus RdRps showing ICTV-accepted virus taxa and other major groups of viruses. Approximate numbers of distinct virus RdRps present in each branch are shown in parentheses. Symbols to the right of the parentheses summarize the presumed virus host spectrum of a lineage. Green dots represent well-supported branches (>0.7) whereas yellow dot corresponds to a weakly supported branch. (B) Genome maps of a representative set of Branch 1 viruses (drawn to scale) showing major, color-coded conserved domains. When a conserved domain comprises only a part of the larger protein, the rest of this protein is shown in light gray. The locations of such domains are approximated (indicated by fuzzy boundaries). CP, capsid protein; MP, movement protein; S3H, superfamily 3 helicase; SJR1 and SJR2, single jelly-roll capsid proteins of type 1 and 2 (see Figure 7).

#### Branch 2: Picornavirus supergroup

The expansive Branch 2 generally corresponds to the previously described “picornavirus supergroup” (Figures 1 and 3) (7, 31). Some of the virus groups that were previously considered peripheral members of this supergroup, such as totiviruses and nodaviruses, were relocated to different branches in the present tree (Branches 4 and 3, respectively), whereas the viruses of the order *Nidovirales* were moved inside Branch 2 from an uncertain position in the tree. Nevertheless, the core of the supergroup remains coherent, suggestive of common ancestry. Within Branch 2, 3 major clades are strongly supported (Figure 3A); however, many of the internal branches are less reliable, so that the relative positions of partitivirus-picobirnavirus, potyvirus-astrovirus and nidovirus clades within Branch 2 remain uncertain.

**Figure.3.**
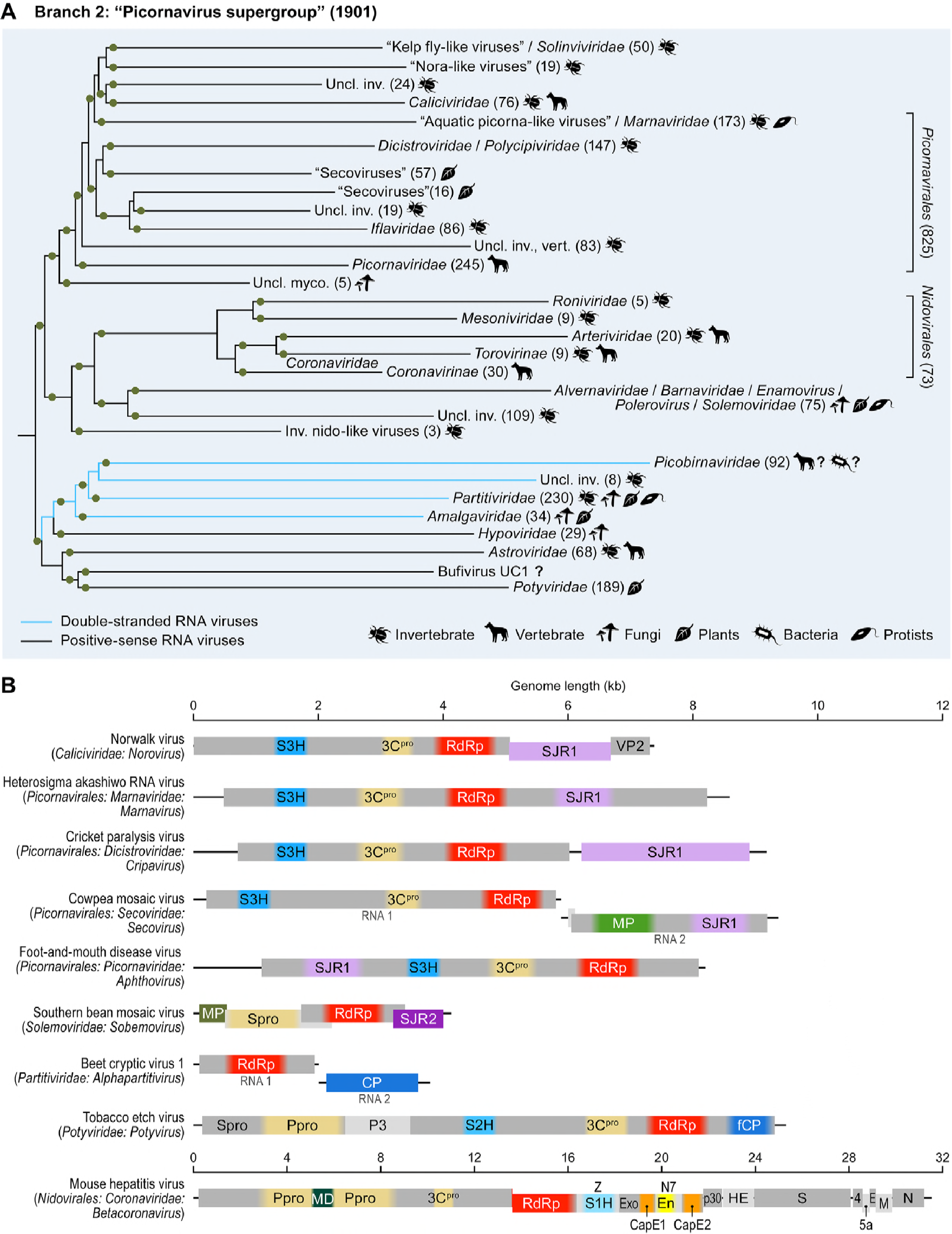
Branch 2 of the RNA virus RNA-dependent RNA polymerases (RdRps): “picornavirus supergroup” of the +RNA viruses expanded to include nidoviruses and two groups of dsRNA viruses, partitiviruses and picobirnaviruses. (A) Phylogenetic tree of the virus RdRps showing ICTV accepted virus taxa and other major groups of viruses. Approximate numbers of distinct virus RdRps present in each branch are shown in parentheses. Symbols to the right of the parentheses summarize the presumed virus host spectrum of a lineage. Green dots represent well-supported branches (>0.7). Inv., viruses of invertebrates (many found in holobionts making host assignment uncertain); myco., mycoviruses; uncl., unclassified; vert., vertebrate. (B) Genome maps of a representative set of Branch 2 viruses (drawn to scale) showing major, color-coded conserved domains. When a conserved domain comprises only a part of the larger protein, the rest of this protein is shown in light gray. The locations of such domains are approximated (indicated by fuzzy boundaries). 3C^pro^, 3C chymotrypsin-like protease; CP, capsid protein; E, envelope protein; En, nidoviral uridylate-specific endoribonuclease (NendoU); Exo, 3’-to-5’ exoribonuclease domain; fCP, capsid protein forming filamentous virions; M, membrane protein; MD macro domain; MP, movement protein; MT, ribose-2-O-methyltransferase domain; N, nucleocapsid protein; N7, guanine-N7-methyltransferase; Ppro, papain-like protease; SJR1 and SJR2, single jelly-roll capsid proteins of type 1 and 2; spike, spike protein; S1H, superfamily 1 helicase; S2H, superfamily 2 helicase; S3H, superfamily 3 helicase; VP2, virion protein 2; Z, Zn-finger domain; Spro, serine protease; P3, protein 3. Distinct hues of same color (e.g., green for MPs) are used to indicate the case when proteins that share analogous function are not homologous.

The largest and most coherent of the Branch 2 clades includes the cornerstone of the picornavirus supergroup, the ~826 viruses-strong order *Picornavirales* (56), expanded with caliciviruses, solinviviruses and a multitude of unclassified viruses infecting invertebrates, vertebrates, fungi, protists and undefined hosts (for viruses discovered by metaviromics) (Figure 3) (11, 14, 17, 57-59). The second largest, deep-branching clade consists of two lineages that include, respectively, +RNA and dsRNA viruses (Figure 3). The +RNA virus lineage combines astroviruses and potyviruses, the evolutionary affinity of which is well recognized (31, 60). The dsRNA lineage includes the members of the families *Amalgaviridae, Hypoviridae, Partitiviridae,* and *Picobirnaviridae,* with each of these families greatly expanded by unclassified affiliates. Finally, the ‘middle’ clade is smaller and less diverse: it encompasses nidoviruses, including the longest of all +RNA virus genomes (61), and solemoviruses with much shorter genomes (Figure 3). Notably, some members of the family *Luteoviridae* and Heterocapsa circularisquama RNA virus, the only known alvernavirus (62), are nested within the solemovirus clade. Given the lack of support beyond the phylogenetic affinity of the RdRps and the dramatic differences in the genomic architectures of nidoviruses and solemoviruses, the possibility that this unification is caused by a tree construction artifact is difficult to rule out (the branch support notwithstanding).

Hypoviruses, a group of fungal capsid-less RNA replicons, have been traditionally viewed as dsRNA viruses. However, comparison of genome architectures and phylogenetic analysis suggested that hypoviruses are derivatives of potyviruses that have lost the capsid protein (63, 64). In the current RdRp tree, hypoviruses cluster with the dsRNA viruses of the partitivirus-picobirnavirus clade rather than with potyviruses (Figure 3). Whether this position is an artifact of tree construction or whether hypoviruses actually share the RdRps with dsRNA viruses is unclear.

The partitivirus-picobirnavirus clade within Branch 2 represents a transition to the *bona fide*dsRNA Baltimore Class (Figure 3). Typical partitiviruses and picobirnaviruses have minimalist genomes that consist of two dsRNA segments encapsidated separately into distinct 120-subunit T=1 capsids (65-68). These genome segments encode, respectively, RdRps and CPs that are clearly homologous between the two families. The CPs of the partitivirus-picobirnavirus clade have been suggested to be distantly related to those of other dsRNA viruses that belong to Branch 4 (33, 69). Notably, this clade also includes some naked RNA replicons that reproduce in algal mitochondria or chloroplasts, use a mitochondrial genetic code and, in terms of lifestyle, resemble “mitoviruses” (14, 70, 71). By analogy, the origin of the partitivirus-picobirnavirus group from an as-yet undiscovered lineage of prokaryotic RNA viruses seems likely. More specifically, this group of dsRNA viruses could have evolved through reassortment of genomic segments encoding, respectively, a +RNA virus RdRp of Branch 2 (possibly a naked RNA replicon) and a dsRNA virus capsid protein related to those of Branch 4 viruses. The most recently evolved branch of partitiviruses is characterized by larger, 4-6-partite genomes, in contrast to mono-or bipartite genomes in the deeper branches (14). This observation emphasizes a major tendency in virus evolution: increase in genome complexity via gradual acquisition of accessory genes (72).

Apart from the SJR-CP, an apparently ancestral protein that is likely to be a shared derived character (synapomorphy) of Branch 2 is a serine protease that is present in members of the order *Picornavirales* (with the diagnostic substitution of cysteine for the catalytic serine), members of the potyvirus-astrovirus clade, solemoviruses, alvernavirus, and nidoviruses (Figure 3B and S2B). As demonstrated previously, this viral protease derives from a distinct bacterial protease, probably of mitochondrial origin, which is compatible with an early origin of Branch 2 in eukaryotic evolution (31).

The reconstruction of protein gain-loss, together with the comparison of genome architectures in this branch, reveal extensive rearrangements as well as gene and module displacement (Figure 3B and S2B). Branch 2 includes viruses with relatively long genomes and complex gene repertoires (nidoviruses, potyviruses and many members of *Picornavirales)* along with viruses with much shorter genomes and minimal sets of genes (astroviruses and solemoviruses). Clearly, evolution of Branch 2 viruses involved multiple gene gains. Of special note is the gain of 3 distinct helicases in 3 clades within this branch: superfamily 3 helicases (S3H) in members of *Picornavirales,* superfamily 2 helicases (S2H) in potyviruses, and superfamily 1 helicases (S1H) in nidoviruses (Figure 3B and S2B). This independent, convergent gain of distinct helicases reflects the trend noticed early in the study of RNA virus evolution, namely, that most viruses with genomes longer than ~6 kb encode helicases, whereas smaller ones do not. This difference conceivably exists because helicase activity is required for the replication of longer RNA genomes (73). Another notable feature is the change of virion morphology among potyviruses (replacement of ancestral SJR-CP by an unrelated CP forming filamentous virions) and nidoviruses (displacement by a distinct nucleocapsid protein). The dramatic change in virion morphology and mode of genome encapsidation might have been necessitated by the inability of SJR-CP-based icosahedral capsids to accommodate the larger genomes of the ancestral potyviruses and nidoviruses. In addition, nidoviruses gained capping enzymes (CapE; Figure 3B and S2B) that most likely were acquired independently of the capping enzymes of other RNA viruses. Nidoviruses also gained ribonucleases and other accessory proteins that are involved in genome replication, virulence and other aspects of the infection cycle of these largest known RNA viruses (61, 74-76).

#### Branch 3: “Alphavirus supergroup”, “flavivirus supergroup” and the extensive diversity of “tombus-like viruses”

Branch 3 is the part of the +RNA virus RdRp tree that underwent the most dramatic rearrangements compared to previous versions. This branch consists of two strongly supported clades of +RNA viruses: i) the assemblage that originally was defined as the “alphavirus supergroup” (7, 15) joined by several additional groups of viruses; and ii) flaviviruses and related viruses (“flavivirus supergroup”; Figure 1 and 4). In the former clade, the alphavirus supergroup encompasses an enormous diversity of plant, fungal, and animal +RNA viruses, and consists of 3 well-supported lineages: tymoviruses, virgaviruses/alphaviruses/endornaviruses, and hepeviruses/benyviruses, each accompanied by related viruses that often form yet unclassified lineages (Figure 4A). Within the “alphavirus supergroup” alone, the genome lengths range from ~6 to ~20 kb. Despite this length variation, all supergroup members harbor a conserved RNA replication gene module encoding a CapE, S1H, and RdRp; the conservation of this module attests to the monophyly of the supergroup (Figure 4B).

**Figure.4.**
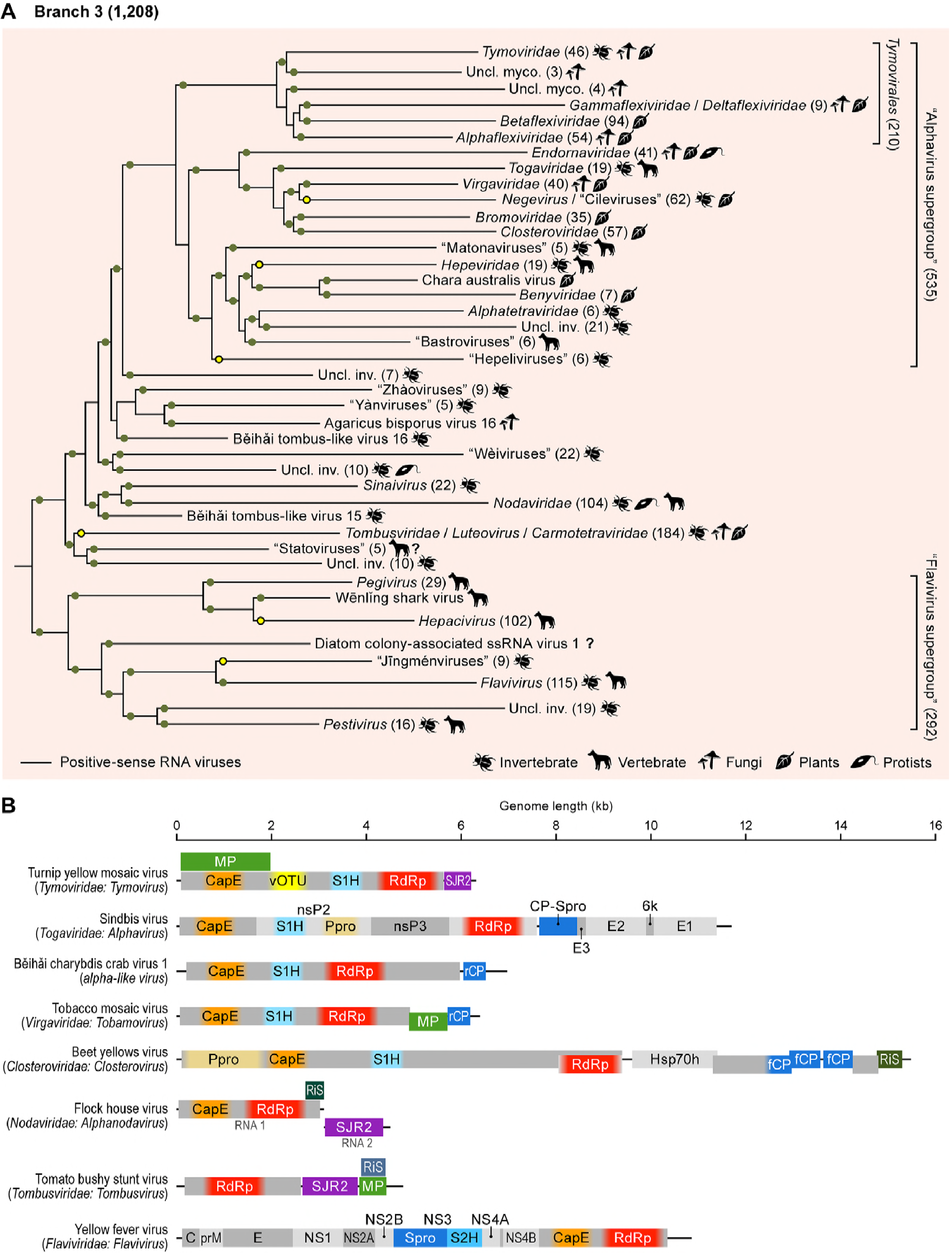
Branch 3 of the RNA virus RNA-dependent RNA polymerases (RdRps): Alphavirus superfamily, radiation of related tombusviruses, nodaviruses and unclassified viruses, and flavivirus supergroup. (A) Phylogenetic tree of the virus RdRps showing ICTV-accepted virus taxa and other major groups of viruses. Approximate numbers of distinct virus RdRps present in each branch are shown in parentheses. Symbols to the right of the parentheses summarize the presumed virus host spectrum of a lineage. Green dots represent well-supported branches (>0.7), whereas yellow dots correspond to weakly supported branches. Inv., viruses of invertebrates (many found in holobionts making host assignment uncertain); myco., mycoviruses; uncl., unclassified. (B) Genome maps of a representative set of Branch 3 viruses (drawn to scale) showing major, color-coded conserved domains. When a conserved domain comprises only a part of the larger protein, the rest of this protein is shown in light gray. The locations of such domains are approximated (indicated by fuzzy boundaries). C, nucleocapsid protein; CapE, capping enzyme; CP-Spro, capsid protein-serine protease; E, envelope protein; fCP, divergent copies of the capsid protein forming filamentous virions; Hsp70h, Hsp70 homolog; MP, movement protein; NS, nonstructural protein; nsP2-3, non-structural proteins; Ppro, papain-like protease; prM, precursor of membrane protein; rCP, capsid protein forming rod-shaped virions; RiS, RNA interference suppressor; S1H, superfamily 1 helicase; S2H, superfamily 2 helicase; SJR2, single jelly-roll capsid proteins of type 2; Spro, serine protease; vOTU, virus OTU-like protease; NS, nonstructural protein. Distinct hues of same color (e.g., green for MPs) are used to indicate the cases when proteins that share analogous function are not homologous.

In contrast, virion architectures vary dramatically even within each of the three lineages of the “alphavirus supergroup”. The major structural themes include: variants of icosahedral capsids formed by SJR-CP (e.g., bromoviruses, tymoviruses); unrelated icosahedral capsids enveloped in a lipoprotein bilayer (togaviruses); flexuous filamentous capsids formed by a distinct type of CP (alphaflexiviruses, betaflexiviruses, gammaflexiviruses, closteroviruses); and rigid rod-shaped capsids assembled from another distinct CP (benyiviruses, virgaviruses). It was traditionally thought that the latter capsid type is specific to viruses of flowering plants (20). However, the recent discovery of a virgavirus-like CP in invertebrate viruses (e.g., Beihai charybdis crab virus 1 in Figure 4B) (14) suggests that the emergence of this unique CP fold antedates land colonization by plants by ~100 Mya. Yet another ‘structural’ theme is offered by endornaviruses, naked RNA replicons which, similarly to hypoviruses in Branch 2 (see above), originally were classified as dsRNA viruses. However, endornaviruses possess all the hallmarks of the alphavirus supergroup and clearly are derived from +RNA viruses of this group. They seem to have been mislabeled dsRNA viruses due to the accumulation of dsRNA replication intermediates in infected cells (18, 77). A parallel loss of the CP genes apparently occurred in deltaflexiviruses which, in RdRp phylogenies, form a sister group to the flexible filamentous gammaflexiviruses (78), and in umbraviruses that are included in the family *Tombusviridae*based on the RdRp phylogeny. Notably, unlike most other capsid-less viruses that are vertically inherited, umbraviruses can hijack capsids of co-infecting luteoviruses for aphid transmission (79).

Within Branch 3, the phylogenetically compact alphavirus supergroup is embedded within the radiation of diverse virus groups including the well-known tombusviruses and nodaviruses, along with several newcomers discovered via metaviromics, such as the “statovirus”, “weivirus”, “yanvirus”, and “zhàovirus” groups (14, 80, 81) (Figure 4A). Our RdRp analysis revealed remarkable phylogenetic heterogeneity within and among these groups and split “tombus-like viruses” into 5 lineages with distinct evolutionary affinities (groups ‘Uncl. inv.’, and subsets of “tombus-like viruses” and “nodaviruses” in Figure 4A). This subdivision is also supported by the analysis of the CPs of these viruses (see section on SJR-CP evolution; Fig. 7). Therefore, in contrast to the alphavirus supergroup, nodaviruses or flavivirus supergroup, the term ‘tombus-like’ loses its evolutionary and taxonomic coherence. Accordingly, we use the term “tombusviruses” (without quotation marks) only for one lineage that includes the members of the current family *Tombusviridae* along with a broad variety of related plant and invertebrate holobiont viruses (14).

The previously suggested, tenuous *Flaviviridae-Tombusviridae* affinity is gone in the present tree although members of both families belong to the same major Branch 3. Plant tombusviruses (and members of closely related plant virus genera), the only group of “tombus-like viruses” that was available at the time of previous analyses (7, 21), now form but a small twig deep within the large assemblage we refer to as tombusviruses. Tombusviruses are affiliated with “statoviruses” (80) and a subset of unclassified viruses from invertebrate holobionts rather than with flaviviruses (Figure 4A). Flaviviruses now form a separate clade within Branch 3, the flavivirus supergroup that includes members of four recognized flaviviral genera *(Pegivirus, Hepacivirus, Flavivirus,* and *Pestivirus),* the newly discovered “jingmenviruses” with segmented genomes (38) and a variety of unclassified, extremely divergent “flavi-like viruses” of animals and plants. This clade is split into two well-supported lineages, one of which includes pegiviruses and hepaciviruses, and the other one consists of the rest of flaviviruses (Figure 4A). Flaviviral virions are enveloped, with the envelope proteins forming an external icosahedral shell, whereas the core nucleocapsid is apparently disordered; the evolutionary provenance of the core protein, with its distinct fold, is unclear (19, 82, 83). Notably, flaviviral envelope proteins are class II fusion proteins that are closely related to alphavirus envelope E1 proteins (84). The theme of gene swapping between these distantly related virus groups of Branch 3 is further emphasized by the homology between alphavirus CPs which form icosahedral capsids under the lipid envelopes, and flavivirus non-structural NS3 proteases that share a chymotrypsin-like fold (84). Because the RdRp tree topology implies that the alphavirus ancestor is more recent than the ancestor of flaviviruses (Figure 1), such adoption of the NS3 protease for a structural role is suggestive of emerging alphaviruses borrowing their structural module from preexisting flaviviruses (19).

The hallmark of Branch 3 is the capping enzyme (CapE), which is present in the entire “alphavirus supergroup” and in flaviviruses (Figure 4 and S2B). A highly divergent version of CapE has been identified in nodaviruses (85) and, in our present analysis, in the additional subset of viruses that grouped with nodaviruses, as well as a few viruses scattered throughout the clade. Formally, CapE is inferred to be ancestral in the entire Branch 3. However, CapEs of “alphavirus supergroup” members, nodaviruses, and flaviviruses are only distantly related to one another, and at least the latter have closer eukaryotic homologs, the FtsJ family methyltransferases (86, 87). Furthermore, tombusviruses, statoviruses, yanviruses, zhaoviruses, weiviruses, and members of the *Pegivirus-Hepacivirus* lineage of flaviviruses lack CapE, putting into question its presence in the ancestor of this branch (Figure 4B, S2B). The most credible evolutionary scenario seems to involve convergent acquisition of CapEs on at least 3 independent occasions, recapitulating the apparent history of helicases in Branch 2 (see above; Figure 3B and S2B). The trend of the capture of helicases by +RNA viruses with longer genomes also holds in Branch 3 and includes the acquisition of S1H at the base of the alphavirus supergroup and S2H by the ancestral flavivirus (Figure 4B, S2B).

To an even greater extent than in Branch 2, the apparent routes of virus evolution in Branch 3 involve lineage-specific gene capture resulting in evolution of complex genome architectures (Figure 4B). The most notable cases are closteroviruses and divergent flaviviruses that have genomes of up to 20-26 kb, rivalling coronaviruses in terms of genome length and the complexity of the gene repertoire (38, 88-90).

The lack of genes assigned to the common ancestor of Branch 3 (with the obvious exception of the RdRp) prevents development of a coherent evolutionary scenario for the entire branch. In the case of the clade encompassing the “alphavirus supergroup” and related viruses, a potential common ancestor could be a simple virus that encoded only an RdRp and a SJR-CP, a CP fold most broadly represented in this clade including diverse tombusviruses, nodaviruses and members of *Bromoviridae,* and *Tymoviridae* within the “alphavirus supergroup”. Proposing such an ancestor for the flavivirus clade is challenged by the lack of viruses with short and simple genomes among flaviviruses. Indeed, the lengths of the genomes in this clade vary from ~9 kb to ~26 kb, with even the shortest ones encoding at least three of the flavivirus signature genes (serine protease [Spro], S2H, and RdRp). One potential clue, however, is provided by “jingmenviruses” with tetra-partite genomes in which the protease-helicase modules and the RdRps are encoded by separate genome segments; two other segments apparently encode structural proteins of unclear provenance (38). This genome architecture could hint at an ancestral flavivirus genome that was assembled from genes borrowed from pre-existing viruses, one of which possessed a divergent “tombus-like virus” RdRp. Although the origins of Branch 3 are murky, major trends in its subsequent evolution clearly included lineage-specific gene capture, starting with helicases and CapEs in the ancestors of the major lineages and followed by diverse genes in smaller groups (Figure 4B).

#### Branch 4: dsRNA viruses

Branch 4, which joins Branch 3 with weak support, includes the bulk of the dsRNA viruses (Figure 1 and 5). All dsRNA viruses in this branch share a unique virion organization and encode homologous CPs. In particular, the specialized icosahedral capsids of these viruses, involved in transcription and replication of the dsRNA genome inside cells, are constructed from 60 homo-or heterodimers of CP subunits organized on an unusual *T=1* (also known as pseudo-7=2) lattice (69, 91). The only exception are the chrysoviruses which encode large CPs corresponding to the CP dimers of other dsRNA viruses and form genuine T=1 capsids (92). Icosahedral capsids of partitiviruses and picobirnaviruses, which encode RdRps belonging to Branch 2, are also constructed from 60 homodimers (66, 67, 93) and have been suggested to be evolutionarily related to those of the dsRNA viruses from RdRp Branch 4 (94) despite little structural similarity between the corresponding CPs. Totiviruses, many of which have “minimal” genomes encoding only RdRps and CPs, comprise one of the two major clades in Branch 4, whereas cystoviruses, the only known prokaryotic dsRNA viruses, together with the vast family *Reoviridae,* which consists of multi-segmented dsRNA viruses infecting diverse eukaryotes, comprise the second clade (Figure 5). The closer phylogenetic affinity between cystoviruses and reoviruses appears to be corroborated by the fact that the inner T=1 icosahedral capsid is uniquely encased by the outer icosahedral shell constructed on a 7=13 lattice in both families (34). Both cystoviruses and reoviruses appear to have gained many clade-specific genes, in particular, RecA-like packaging ATPases of the former (95) and the CapEs of the latter that are only distantly related to CapEs of other RNA viruses and likely were acquired independently (96, 97) (Figure 5, S2B).

**Figure.5.**
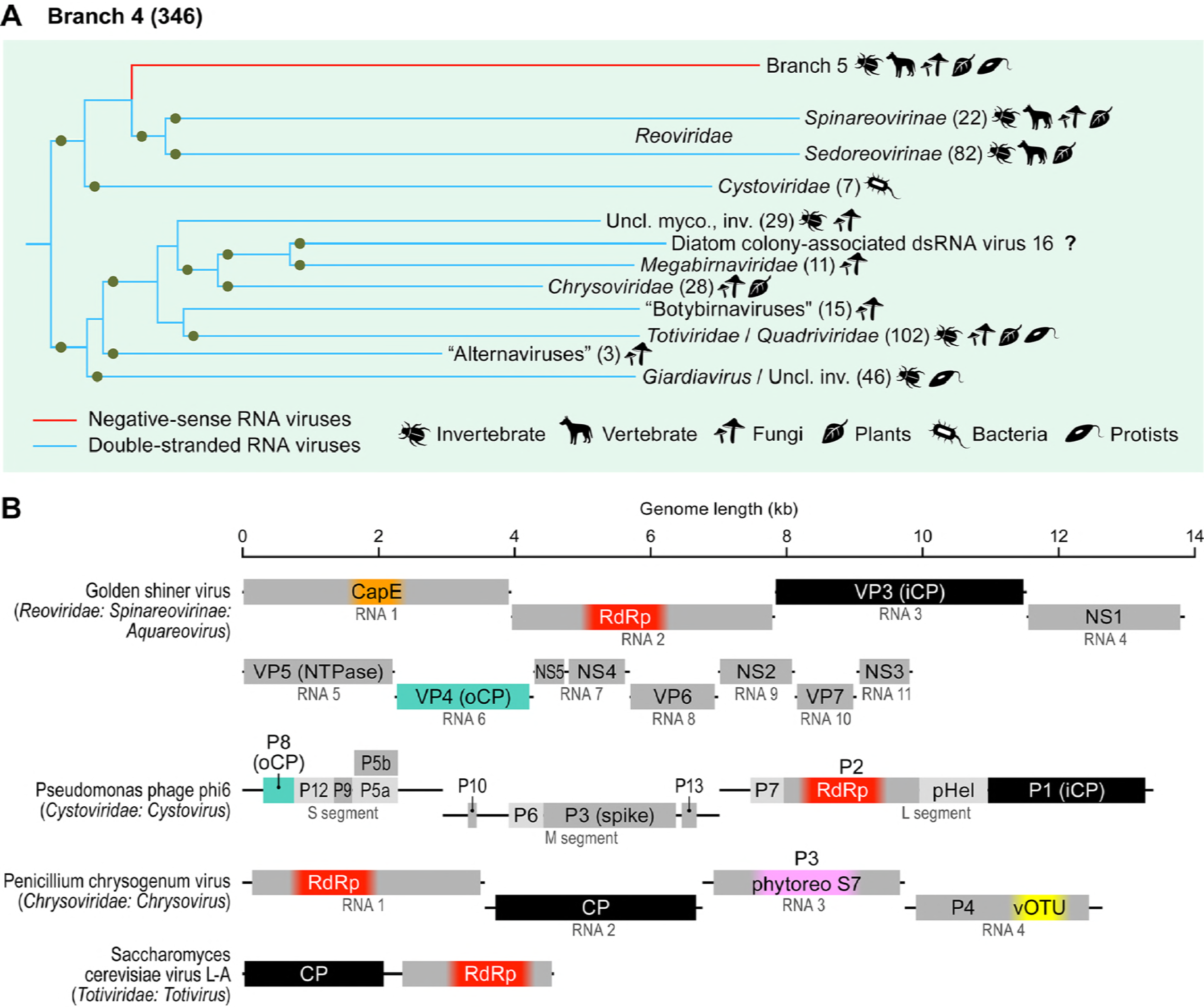
Branch 4 of the RNA virus RNA-dependent RNA polymerases (RdRps): dsRNA viruses of eukaryotes and prokaryotes. (A) Phylogenetic tree of the virus RdRps showing ICTV-accepted virus taxa and other major groups of viruses. Approximate numbers of distinct virus RdRps present in each branch are shown in parentheses. Symbols to the right of the parentheses summarize the presumed virus host spectrum of a lineage. Inv., viruses of invertebrates (many found in holobionts making host assignment uncertain); myco., mycoviruses; uncl., unclassified. Green dots represent well-supported branches (>0.7). (B) Genome maps of a representative set of Branch 4 viruses (drawn to scale) showing major, color-coded conserved domains. When a conserved domain comprises only a part of the larger protein, the rest of this protein is shown in light gray. The locations of such domains are approximated (indicated by fuzzy boundaries). CapE, capping enzyme; CP, capsid protein; iCP, internal capsid protein; NS, non-structural protein; NTPase, nucleotide triphosphatase; oCP, outer capsid protein; P, protein; phytoreoS7, homolog of S7 domain of phytoreoviruses; pHel, packaging helicase; vOTU, virus OTU-like protease; VP, viral protein; The CPs of totiviruses and chrysoviruses are homologous to iCPs of reoviruses and cystoviruses (black rectangles).

#### Branch 5: -RNA viruses

Branch 5, the 100% supported lineage combining all-RNA viruses, is lodged within Branch 4 as the sister group of reoviruses, and this position is upheld by two strongly supported internal branches in the RdRp tree (Figure 1 and 6). The-RNA branch splits into 2 strongly supported clades. The first clade encompasses the 348 viruses-strong membership of the order *Mononegavirales* (98), along with the members of the distantly related family *Aspiviridae* (99), 3 groups of-RNA viruses discovered through metaviromics (“chuviruses”, “qínviruses”, “yuèviruses”) (14, 39), and a group of unclassified fungal viruses (Figure 6A) (42, 100). In contrast to the members of the *Mononegavirales,* most of which possess unsegmented genomes, the remainder of this clade is characterized by bi-, tri-, or even tetrasegmented genomes (Figure 6B). The second clade combines the family *Orthomyxoviridae,*the genus *Tilapinevirus* (101) and the large order *Bunyavirales* (394 viruses) (102). The latter order consists of two branches, one of which is the sister group to the orthomyxovirus/tilapinevirus clade (albeit with weak support). The number of negative-sense or ambisense genome segments in this clade varies from 2-3 in most of bunyaviruses to 8 in viruses of the genus *Emaravirus* and the orthomyxovirus/tilapinevirus group (10, 99, 101, 103). A notable acquisition in the first clade is a CapE, whereas members of the second clade share “cap-snatching” endonucleases (En) (10).

**Figure.6.**
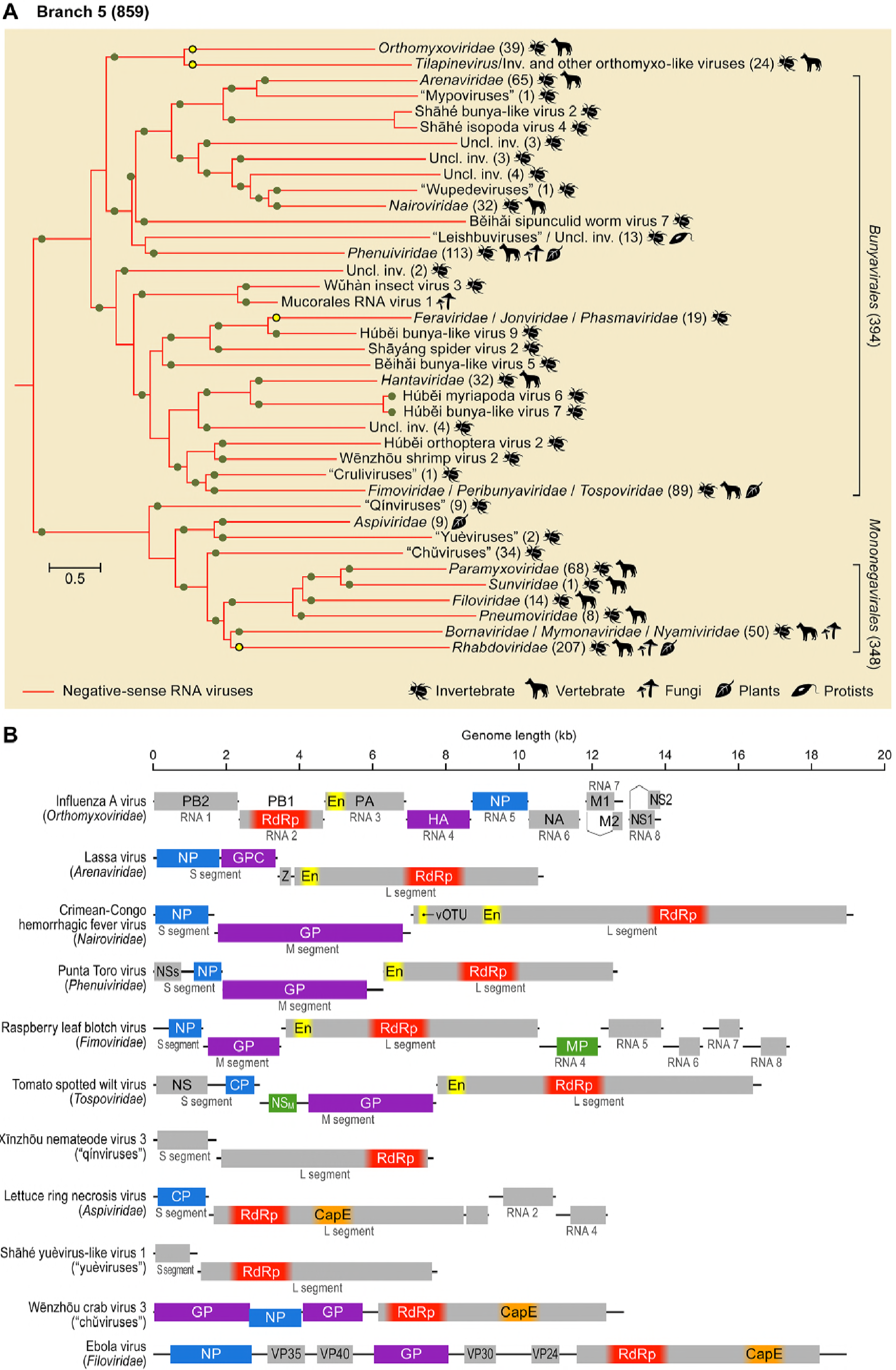
Branch 5 of the RNA virus RNA-dependent RNA polymerases (RdRps):-RNA viruses. (A) Phylogenetic tree of the virus RdRps showing ICTV-accepted virus taxa and other major groups of viruses. Approximate numbers of distinct virus RdRps present in each branch are shown in parentheses. Symbols to the right of the parentheses summarize the presumed virus host spectrum of a lineage. Green dots represent well-supported branches (>0.7), whereas yellow dots correspond to weakly supported branches. Inv., viruses of invertebrates (many found in holobionts making host assignment uncertain); uncl., unclassified. (B) Genome maps of a representative set of Branch 5 viruses (drawn to scale) showing major, color-coded conserved domains. When a conserved domain comprises only a part of the larger protein, the rest of this protein is shown in light gray. The locations of such domains are approximated (indicated by fuzzy boundaries). CapE, capping enzyme; CP, capsid protein; EN, “cap-snatching” endonuclease; GP, glycoprotein; GPC; glycoprotein precursor; HA; hemagglutinin; M, matrix protein; MP, movement protein; NA; neuraminidase; NP, nucleoprotein; NS, nonstructural protein; NS_M_, medium nonstructural protein; NSs, small nonstructural protein; PA, polymerase acidic protein; PB, polymerase basic protein; vOTU; virus OTU-like protease; VP; viral protein; Z, zinc finger protein.

## Patterns of the single jelly-roll capsid protein evolution

The SJR-CP is the dominant type of CP among +RNA viruses and is also found in members of one family of dsRNA viruses (*Birnaviridae*). Structural comparisons indicate that SJR-CPs of RNA viruses form a monophyletic group and likely have been recruited from cellular SJR proteins on a single occasion during the evolution of RNA viruses (19). The short length and high divergence of SJR-CPs preclude adequate resolution in phylogenetic analysis; thus, we performed profile-profile sequence comparisons and clustering of all viral SJR-CP sequences in our dataset (see Methods for details). Analysis of the resulting network revealed patterns that are generally congruent with the RdRp phylogeny and provide further insights into the evolution of different branches of RNA viruses.

At conservative P value thresholds (P<1e^−10^), the majority of SJR-CPs segregated into two large clusters both of which contained representatives from RdRp Branch 2. Cluster 1 included the members of *Picornavirales, Caliciviridae,* and diverse “picorna-like viruses” of invertebrates, whereas cluster 2 consisted of the members of the families *Astroviridae, Luteoviridae*, and *Solemoviridae* and “solemo-like viruses” (Figure S3). In addition, cluster 2 contained members of several families from RdRp Branch 3, namely, *Tombusviridae* (and diverse “tombus-like viruses”), *Hepeviridae,* a subgroup of *Nodaviridae* and “statoviruses”.

At less restrictive P value thresholds (P<1e^−03^), all SJR-CPs were interconnected, largely, through making contacts to the core of cluster 2. Only “ourmiaviruses” had stronger affinity to picornaviruses in cluster 1 (Figure 7). This pattern of connectivity is consistent with the radiation of SJR-CPs from a common ancestor, likely resembling sequences from cluster 2 of Branch 2. This analysis also revealed high CP sequence divergence among members of some families (e.g., *Bromoviridae*) and numerous cases of apparent CP gene replacement. For instance, the CPs of nodaviruses fall into two groups: one is related to the turreted CPs of tetraviruses and the other is similar to CPs of tombusviruses, mirroring the RdRp phylogeny (Figure 7). At a greater phylogenetic distance, CPs of astroviruses and hepeviruses are closely related despite them being affiliated to Branches 2 and 3, respectively, suggesting CP gene replacement in the ancestor of one of the two families. Given that the CPs of hepeviruses connect to SJR-CPs of other viruses through astroviruses, CP gene replacement most likely occurred in the ancestor of hepeviruses (Figure S3). Notably, the CPs of “zhàoviruses”, “wèiviruses”, and “tombus-like” and “solemo-like viruses” (diverse virus assemblage within solemovirus branch, to the exclusion of bona fide *Solemoviridae)* did not form discrete clusters but rather were affiliated with diverse virus groups, suggesting extensive recombination in these viruses, with multiple CP gene exchanges (Figure 7). In the case of unclassified “narnaviruses” and “ourmiaviruses”, the CP genes apparently have been acquired on more than 3 independent occasions from different groups of viruses, emphasizing the impact of recombination and gene shuffling in the evolution of RNA viruses. Previously, a similar extent of chimerism has been also observed among ssDNA viruses (104, 105), highlighting the evolutionary and functional plasticity of short viral genomes.

**Figure.7.**
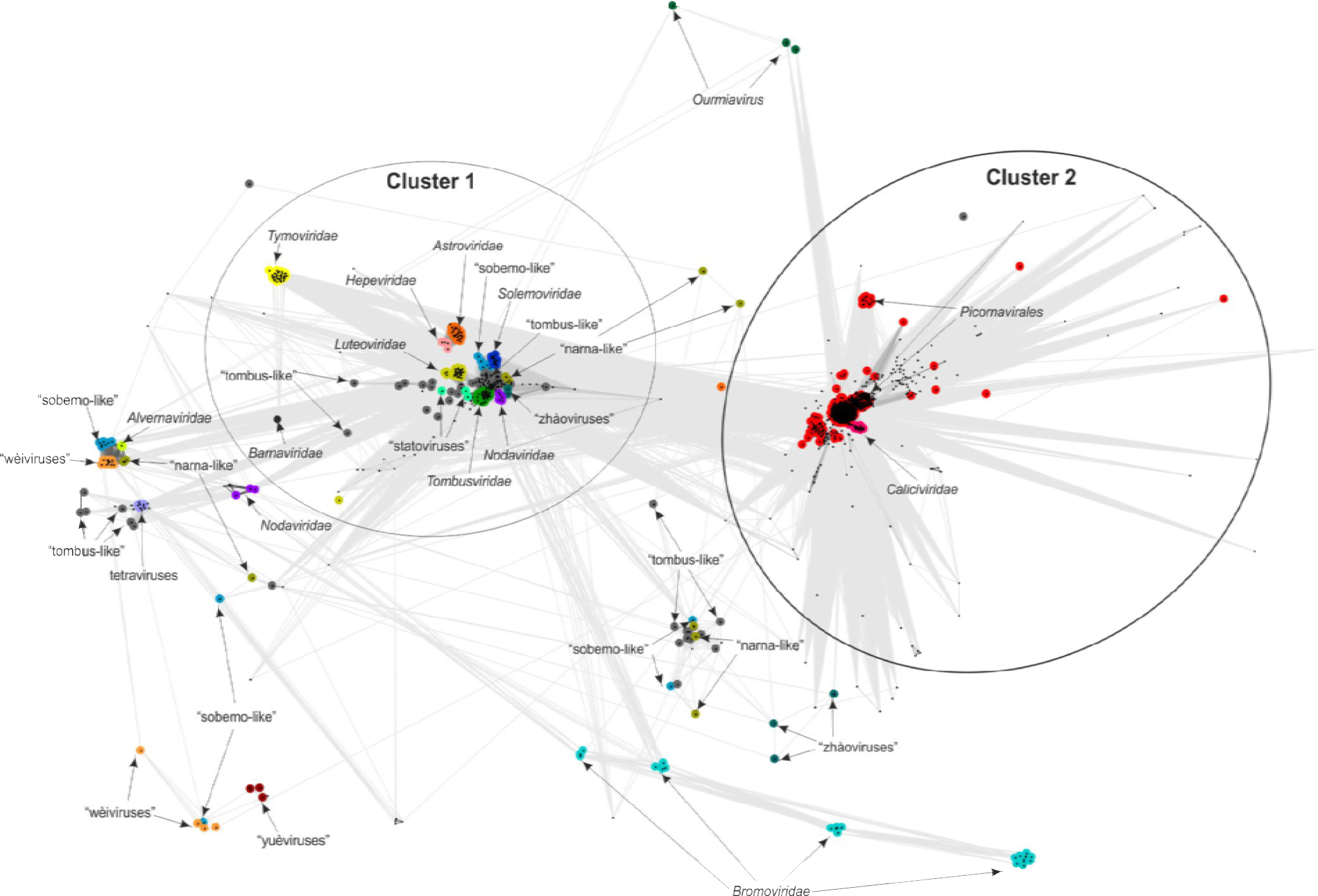
Sequence similarity networks of SJR-CPs. Protein sequences were clustered by the pairwise similarity of their hmm profiles using CLANS. Different groups of SJR-CPs are shown as clouds of differentially colored circles, with the corresponding subgroups labeled as indicated in the figure. Edges connect sequences with CLANS P-value of < 1e-03.

## The modular gene-sharing network of RNA viruses: gene transfer and module shuffling

The pronounced structural and functional modularity of virus proteomes and pervasive shuffling of the genomic regions encoding distinct protein modules are key features of virus evolution (11,15, 17). Therefore, a productive approach to the study of the virosphere that complements phylogenetics is the construction and analysis of networks of gene sharing. Bipartite networks, in which one type of nodes corresponds to genes and the other one to genomes, have been employed to investigate the dsDNA domain of the virosphere (45). This analysis revealed hierarchical modular organization of the network, with several modules including non-obvious connections between disparate groups of viruses (44, 106). Although, for RNA viruses, this type of analysis is less informative due to the small number of proteins encoded in each viral genome, the “pan-proteome” of RNA viruses is large (Data set S5), prompting us to experiment with bipartite gene sharing networks for RNA viruses. The initial search for statistically significant modularity identified 54 distinct modules, most of which included a single virus family (Figure 8A, B). Remarkably, the family *Reoviridae* has been split into 5 modules, highlighting the vast diversity of this family, comparable to that in order-level taxa. Among the exceptions, the most expansive module included the viruses of the order *Picornavirales* (module 29), together with the family *Caliciviridae* (module 47), that are linked through the conserved suit of genes including SJR-CP, chymotrypsin-like protease, S3H and the RdRp (Figure 8A and C). Viruses of the order *Bunyavirales* were also recovered in a single module that is characterized by the presence of a conserved nucleocapsid (with the exception of the families *Nairoviridae* and *Arenaviridae)* and the cap-snatching endonuclease (module 51; Figure 8A and C).

**Figure.8.**
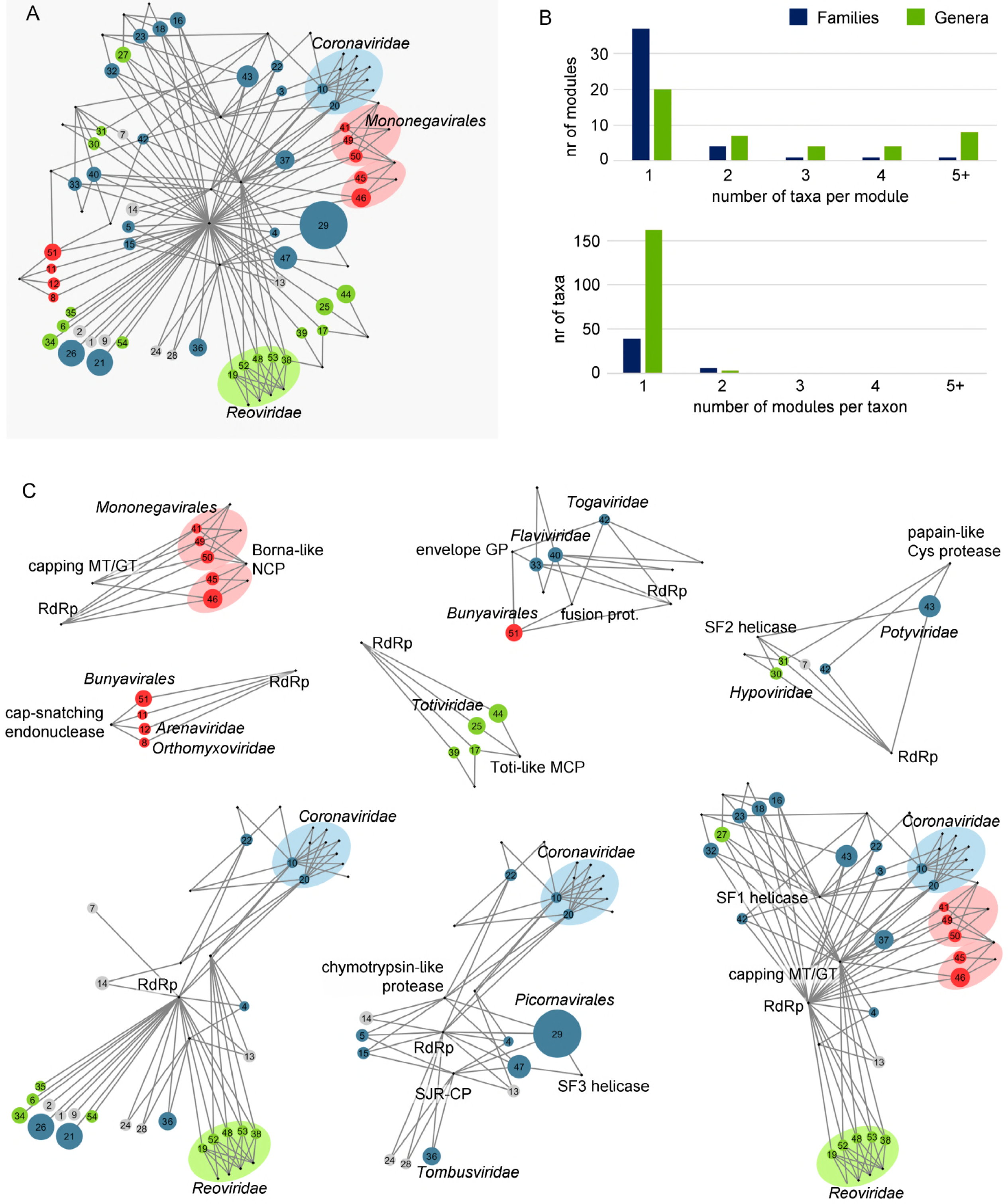
Bipartite network of gene sharing in RNA viruses. (A) Groups of related viruses were identified as the modules of the bipartite genome-gene network (not shown), whereas connector genes were defined as those genes present in two or more modules with prevalence greater than 65%. The network in A shows viral modules as colored circles (blue, +RNA viruses; green, dsRNA viruses; red,-RNA viruses), linked to the connector genes (black dots) that are present in each module. The size of the circles is proportional to the number of genomes in each module. Shaded ovals indicate statistically significant, 1st-order supermodules that join modules from taxonomically related groups. (B) Taxonomic analysis of the network modules confirms that most modules contain viruses from a single family and that families do not tend to split among modules. (C) High-order supermodules of the RNA virus network, obtained by iteratively applying a community detection algorithm on the bipartite network of (super)modules and connector genes. GP, glycoprotein; GT, guanylyltransferase; MT, methyltransferase; NCP, nucleocapsid; RdRp, RNA-dependent RNA polymerase; SF, superfamily; SJR-CP, single jelly roll capsid protein.

The next stage of the network analysis aims at detecting supermodules that are formed from the primary modules via connecting genes. The supermodules of RNA viruses failed to attain statistical significance due to the small number of shared genes but nevertheless, some notable connections are revealed by this analysis. Specifically, 8 overlapping supermodules were identified (Figure 8C). The largest and, arguably, most remarkable is a supermodule that combines +RNA, dsRNA and-RNA viruses that share the capping enzymes, S1H (with the exception of *Reoviridae* and *Mononegavirales)* and additional connector genes (e.g., the OTU family protease) that link some of the constituent modules. The second largest supermodule combines large subsets of viruses from the RdRp Branches 2 and 3 that are connected through the SJR-CP and the chymotrypsin-like protease. Another supermodule encompasses enveloped +RNA viruses of the families *Flaviviridae* and *Togaviridae,* and-RNA viruses of the order *Bunyavirales* (except for *Arenaviridae)* that share homologous envelope glycoproteins (except for flaviviruses) and class II fusion proteins.

Information from gene sharing is inherently limited for RNA viruses due to the small number of genes in each genome. Nevertheless, the bipartite network analysis reveals prominent “horizontal” connections that are underlain either by actual gene exchange or by parallel acquisition of homologous genes by distinct RNA viruses.

## Host ranges of RNA viruses: evolutionary implications and horizontal virus transfer

RNA viruses have been identified in representatives of all major divisions of eukaryotes, whereas in prokaryotes, members of two families of RNA viruses are known to infect only a limited range of hosts (11, 13, 15, 31). For Branch 1 in our phylogenetic tree of RdRps, the route of evolution from leviviruses infecting prokaryotes to eukaryotic “ourmiaviruses” of plants and invertebrates is readily traceable and involves a merger between a levivirus-derived naked RNA replicon that eukaryotes most likely inherited from the mitochondrial endosymbiont with the SJR-CP of a eukaryotic “picorna-like virus”. Notably, such a merger seems to have occurred on at least three other independent occasions in Branch 1 because several groups of invertebrate holobiont “narnaviruses” and some “ourmiaviruses” encode distantly related SJR-CPs that apparently were acquired from different groups of plant and animal viruses (Figure 7).

The case of cystoviruses is less clear given that this clade is sandwiched between eukaryotic viruses in Branch 4 and therefore does not seem to be a good candidate for the ancestor of this branch. It appears more likely that the ancestor was a toti-like virus, whereas cystoviruses are derived forms, which implies virus transfer from eukaryotes to prokaryotes. However, an alternative scenario might be considered. No known prokaryotic viruses are classified in Branch 2 but it has been proposed that picobirnaviruses, for which no hosts have been reliably identified, actually are prokaryotic viruses. This proposal is based on the conspicuous conservation of functional, bacterial-type, ribosome-binding sites (Shine-Dalgarno sequences) in picobirnavirus genomes (107, 108). Should that be the case, viruses of prokaryotes might be lurking among totiviruses as well. Then, Branch 4 would stem from a prokaryotic ancestor avoiding the need to invoke virus transfer from eukaryotes to prokaryotes to explain the origin of the cystoviruses.

We made an attempt to quantify the potential horizontal virus transfer (HVT) events in RNA viruses that represent the 5 major branches of the RdRp tree. The leaves of the tree were labeled with the known hosts, the entropy of the host ranges for each subtree was calculated, and the resulting values were plotted against the distance from the root (Figure 9). By design, for all branches, entropy (host diversity) drops from the maximum values at the root to zero at the leaves. All branches show substantial host range diversity such that, for example, at half-distance from root to leaves, all branches, except for Branch 1, retain at least half of the diversity (Figure 9). Furthermore, differences between the branches are substantial, with the highest entropy observed in branches 4 (dsRNA viruses) and 5 (-RNA viruses). With all the caveats due to potential errors and ambiguities in host assignment, this analysis strongly suggests that HVT played an important role in the evolution of all major groups of RNA viruses.

**Figure.9.**
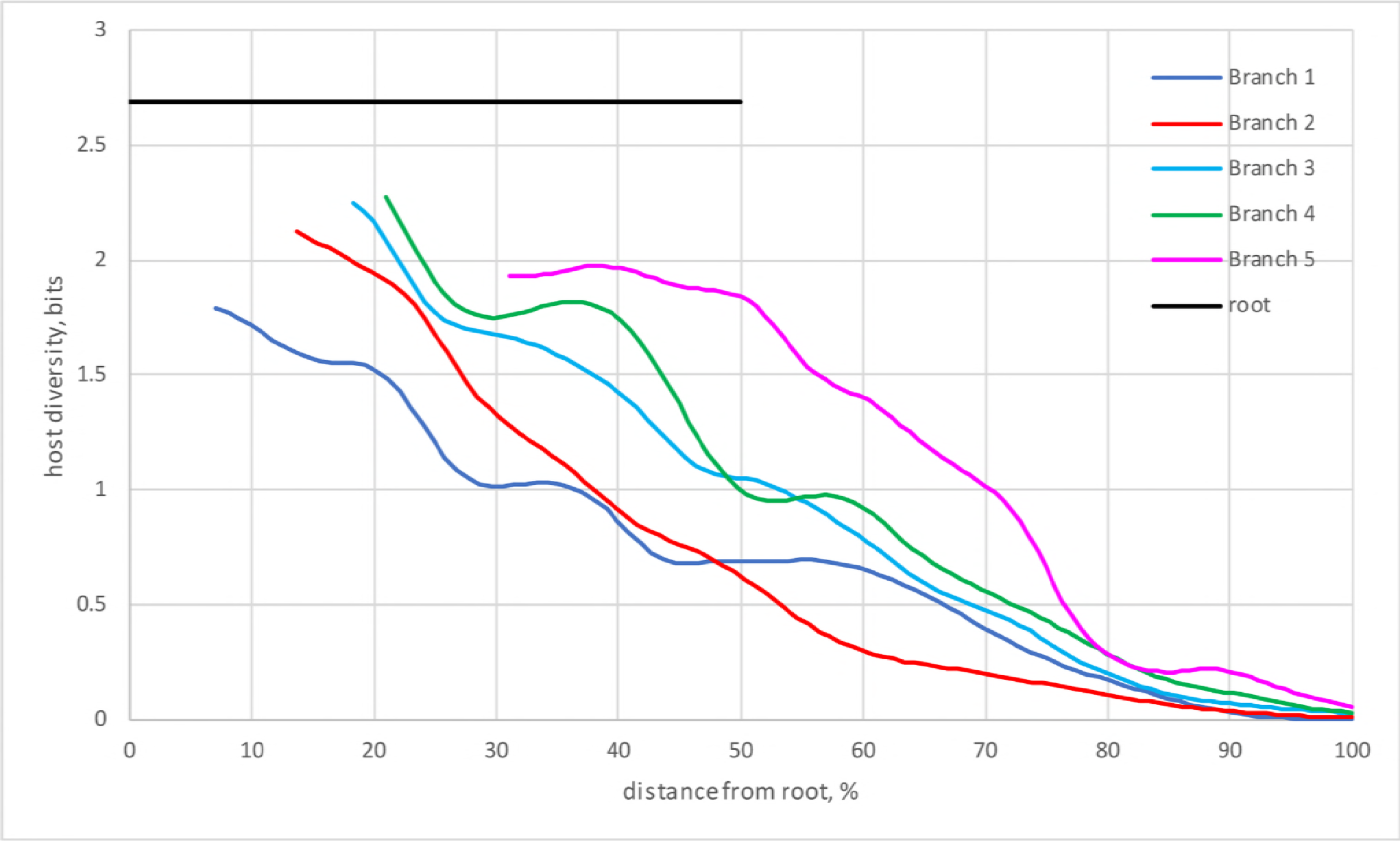
Quantitative analysis of the host range diversity of RNA viruses. The entropy of host ranges is plotted against the ultrameterized tree depth for the 5 main branches of the RdRp phylogeny (see Figure 1).

## DISCUSSION

### RNA virus evolution coming into focus

This work was prompted by the advances of metaviromics, which have dramatically increased the known diversity of RNA viruses (11, 13, 17, 37, 57). We reasoned that this expansion of the RNA virosphere could provide for an improved understanding of virus evolution. Although further progress of metaviromics and enhanced phylogenomic methods will undoubtedly change current ideas, we believe that some key aspects of RNA virus evolution are indeed coming into focus.

The expanded diversity of RNA viruses combined with the iterative procedure for phylogenetic analysis allowed us to obtain a tree of all RdRps and the most closely related RTs in which the main branches are strongly supported and thus appear to be reliable (Figure 1). To our knowledge, the picture of RNA virus evolution emerging from the tree has not been presented previously. The tree seems to clarify the relationships between the 3 Baltimore Classes of RNA viruses by revealing the nested tree structure in which dsRNA viruses evolved, on at least two occasions, from +RNA viruses, whereas -RNA viruses evolved from a distinct group of dsRNA viruses.

The derivation of-RNA viruses from dsRNA viruses is, arguably, the most unexpected outcome of the present analysis, considering the lack of genes (other than the RdRp) shared by these virus classes. Clearly, given that the primary evidence behind the derivation of-RNA viruses from within dsRNA viruses comes from deep phylogeny, extreme caution is due in the interpretation of this observation. However, the pronounced similarity between the 3D structures of the RdRps of the-RNA influenza virus A and bacteriophage ^6 dsRNA cystovirus (35) is compatible with our findings. Further, because virtually no-RNA viruses are known in prokaryotes or unicellular eukaryotes [with the single exception of “leischbuviruses” in parasitic trypanosomatids (50) that were likely acquired from the animal hosts of these protists], their later origin from a preexisting group of +RNA or dsRNA viruses appears most likely.

The +RNA to dsRNA to-RNA scenario of RNA virus genome evolution also makes sense in terms of the molecular logic of genome replication-expression strategies. Indeed, +RNA viruses use the simplest genomic strategy and, in all likelihood, represent the primary pool of RNA viruses. The dsRNA viruses, conceivably, evolved when a +RNA virus switched to encapsidating a replicative intermediate (dsRNA) together with the RdRp. Naked replicons similar to “mitoviruses”, hypoviruses, and endornaviruses might have been evolutionary intermediates in this process. This switch does not seem to be as “easy” and common as previously suspected (15, 32) but, nevertheless, appears to have occurred at least twice during the evolution of RNA viruses. The origin of-RNA viruses is the next step during which the plus-strand is discarded from the virions, perhaps simplifying the processes of transcription and replication. Conceivably, the evolution of dsRNA and-RNA viruses, in which transcription and replication of the viral genomes are confined to the interior of virions or nucleocapsid transcription/replication complexes and no dsRNA accumulates in the infected cells, was driven by the advantage of escaping some of the host defense mechanisms, in particular, RNA interference (109, 110). The membrane-associated replication complexes of +RNA viruses could represent an initial step in this direction (9).

Obviously, evolution of the RdRp does not equal evolution of viruses: other genes, in particular those encoding capsid and other structural proteins, are crucial for virus reproduction, and these genes often have different histories. The reconstruction of gene gain and loss sheds some light on these aspects of RNA virus evolution. The ancestors of each of the major branches of RNA viruses except for Branch 1 appear to have been simply organized +RNA viruses resembling tombusviruses (Figure 10). Thus, these types of viruses encoding RdRps and SJR-CPs might have been ancestral to the bulk of eukaryotic RNA viruses (apart from those in Branch 1 that derive directly from prokaryotic leviviruses). The subsequent parallel capture of different helicases enabled evolution of increasingly complex genomes via accumulation of additional genes (Figure 10). Notably, similar levels of complexity, with the complete coding capacity of 20 to 40 kb, were reached independently in 4 branches of the RdRp tree, namely, Branch 2 (coronaviruses), Branch 3 (closteroviruses and flaviviruses), Branch 4 (reoviruses), and Branch 5 (filoviruses and paramyxoviruses) (Figure 3-6).

**Figure.10.**
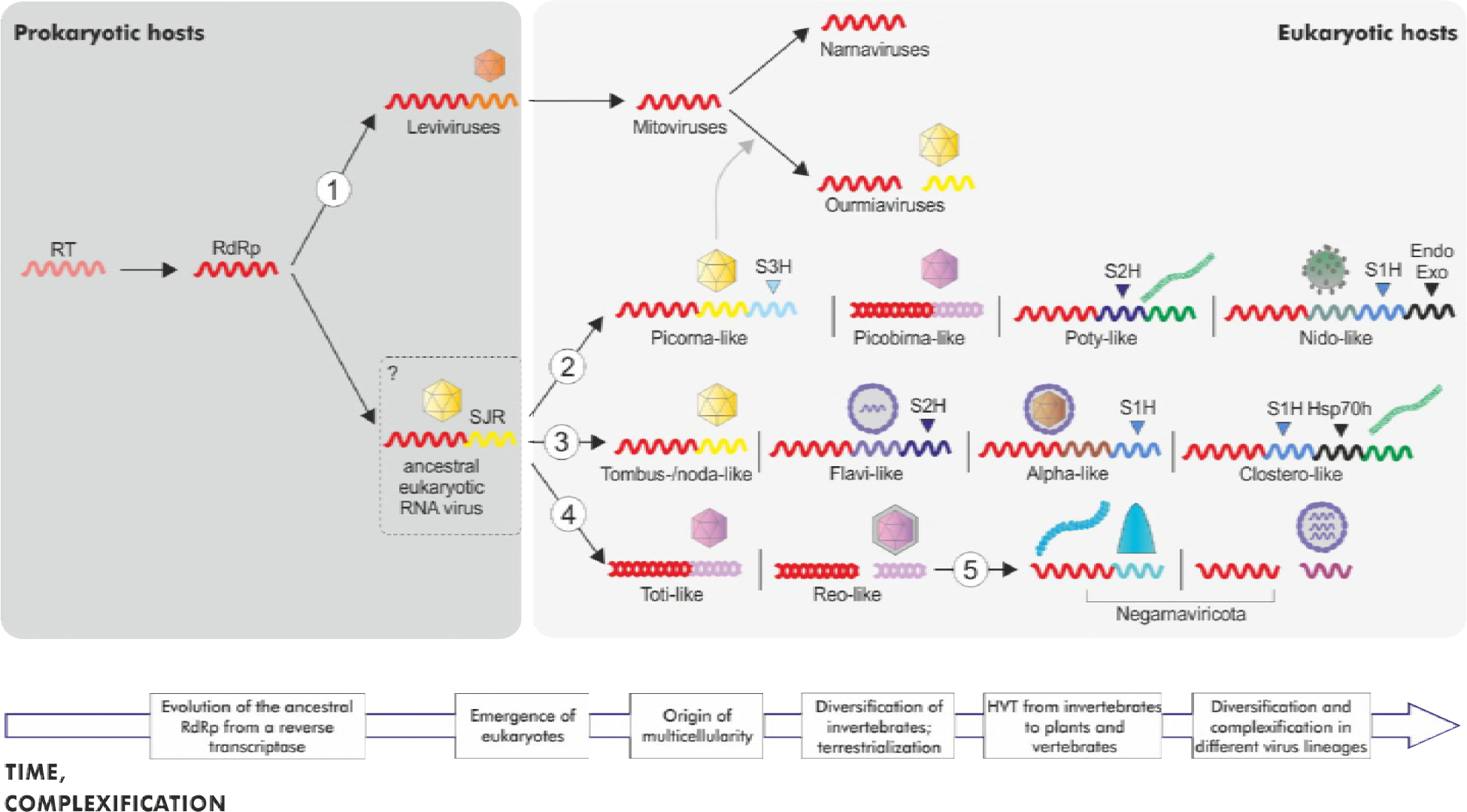
A general scenario of RNA virus evolution. The figure is a rough scheme of the key steps of RNA virus evolution inferred in this work. The main branches from the phylogenetic tree of the RdRps are denoted 1 to 5 as in Figure 1. Only the RdRp, CP genes and helicase (S1H, S2H and S3H for the helicases of superfamilies 1, 2 and 3, respectively) are shown systematically. The helicases appear to have been captured independently and in parallel in 3 branches of +RNA viruses, facilitating the evolution of larger, more complex genomes. Additional genes, namely, Endo and Exo (for endonuclease and exonuclease, respectively) and Hsp70h (heat shock protein 70 homolog), are shown selectively, to emphasize the increased genome complexity, respectively, in *Nidovirales* and in *Closteroviridae.* The virion architecture is shown schematically for each included group of viruses. Icosahedral capsids composed of unelated CPs are shown by different colors (see text for details). The question mark at the hypothetical ancestral eukaryotic RNA virus indicates the uncertainty with regard to the nature of the host (prokaryotic or eukaryotic) of this ancestral form. The block arrow at the bottom shows the time flow and the complexification trend in RNA virus evolution.

Gene exchange and shuffling of gene modules are important factors of RNA virus evolution. For example, it appears that all dsRNA viruses in the two major clades within Branches 2 and 4 share homologous structural modules that combine with distinct RdRps. At least in the case of partiti-picobirnaviruses in Branch 2, the dsRNA virus (“toti-like virus”) CP apparently displaced the ancestral SJR-CP. However, this particular protein structure does not seem to be essential to encapsidate a dsRNA genome: birnaviruses, whose provenance is uncertain due to the permutation in their RdRps, retain SJR-CP, which is most closely related to SJR-CP of nodaviruses and tetraviruses (19). An even more striking example of module shuffling is presented by amalgaviruses, dsRNA viruses that group with partitiviruses in the RdRp tree but encode a distant homolog of the nucleocapsid protein of-RNA bunyaviruses (111-114). More generally, structural and replication modules have been repeatedly shuffled during the evolution of +RNA viruses. Examples include displacement of the ancestral SJR-CP by a filamentous CP in potyviruses and by a helical nucleocapsid protein in nidoviruses, and multiple cases of displacement with rod-shaped-like CP and unique nucleocapsid proteins in Branch 3. Thus, exchange of genes and gene modules among RNA viruses is pervasive and can cross the boundaries of Baltimore Classes.

Another recurring trend in RNA virus evolution is the loss of the structural module resulting in the emergence of naked RNA replicons such as “narnaviruses” and “mitoviruses” in Branch 1, hypoviruses in Branch 2, and endornaviruses, umbraviruses and deltaflexiviruses in Branch 3 (18). On some occasions, broad horizontal spread of a gene leads to a major shift in the lifestyle of viruses, such as adaptation of viruses to a new type of hosts. The primary cases in point are the movement proteins (MPs) of plant viruses that are represented in all 5 branches of the RdRp tree and, outside of the RNA part of the virosphere, in plant caulimoviruses and badnaviruses, and ssDNA viruses (115).

### The prevalence of HVT and the overall course of RNA virus evolution

Arguably, the most striking realization brought about by metaviromics is the diverse host range of numerous groups of viruses, even tight ones that occupy positions near the tips of the RdRp tree. Extending early observations on then highly unexpected similarities between viruses of animals and plants, recent metaviromic analyses reveal numerous clusters of indisputably related viruses infecting animals and plants, plants and fungi, and in some cases, animals or plants and protists. The evolutionary relationships between viruses with distinct host ranges are supported not only by the phylogeny of the RdRp, but also by the fact that these viruses share additional conserved domains, such as, for example, SJR1, S3H and 3C-Pro in the case of *Picornavirales*members infecting protists, plants, fungi, invertebrates and vertebrates (Fig. 3).

Invertebrates are particularly promiscuous hosts for viruses, often sharing the same virus group with distantly related organisms. Certainly, much caution is due in the interpretation of host range assignments from metaviromics, especially, holobiont studies. Viruses identified in holobiont samples of, say, invertebrates could actually infect protists associated with these animals (50) or could represent contamination from fungal, plant or even prokaryotic sources. These uncertainties notwithstanding, the extensive diversity of hosts even within small branches of the RdRp tree is undeniable. A coevolution scenario in which the ancestors of all these viruses originated from the common ancestor of the respective groups of eukaryotes and coevolved with the hosts implies an enormous diversity of RNA viruses in early eukaryotes. This scenario appears to be highly unlikely given the apparent paucity of RNA viruses in the extant protists (although new metaviromic studies might substantially expand the range of protist viruses). The pervasive HVT alternative seems much more plausible, especially given that arthropods, nematodes, and other invertebrates are well known as virus vectors and thus fit the role of RNA virus reservoirs (11).

In addition to invertebrates that appear to be dominant HVT agents, fungi could also play an important role in HVT within the global RNA virome. Indeed, fungi that are tightly associated with plants and insects often share closely related viruses with these organisms (116-118). Furthermore, an indisputable case of cross-kingdom transfer of an insect iflavirus to an entomopathogenic fungus has been recently described (119).

These findings appear to be best compatible with a grand evolutionary scenario (Figure 10) in which the ancestor of the eukaryotic RNA virome was a levi/narnavirus-like naked RNA replicon that originally reproduced in mitochondria and combined with a host carbohydrate-binding SJR protein (19) or a preexisting SJR-CP from a DNA virus to form a simple ancestral virus. Given that viruses of Branches 2, 3, and 4 are present in modern protists, it appears likely that these branches emerged in early eukaryotes. However, because of the apparent dominance of the viruses of the RdRp Branch 2 (“picornavirus supergroup” in general and “aquatic picorna-like viruses” clade in particular) in protists, this branch likely diversified first, whereas the diversification of Branches 3 and 4 occurred later, after ancestral protist viruses were transferred to marine invertebrates during the Cambrian explosion. The recent analysis of the viromes of ctenophores, sponges and cnidarians suggests that substantial diversification of RNA viruses occurred already in these deeply branching metazoa (120). Invertebrates brought their already highly diverse RNA virome to land at terrestrialization and subsequently inoculated land plants. In land plants, RNA viruses, particularly those of Branch 3, dramatically expanded, in part, perhaps, because of the exclusion of competing large DNA viruses. Finally, it seems plausible that, given the high prevalence of-RNA viruses in metazoa and their virtual absence in protists [with the exception of the recently discovered “leishbuviruses” that likely invaded their parasitic protist hosts via HVT from an animal host (49, 50)], these viruses that comprise the RdRp Branch 5 evolved in animals via mixing and matching genes from reovirus-like and flavivirus-like ancestors.

### The impending overhaul of RNA virus taxonomy

The expansion of the global RNA virome thanks to the advances of metaviromics, combined with the phylogenomics results, seem to call for an overhaul of the current virus taxonomy on multiple levels. Most importantly, creation of a coherent, hierarchical system with multiple taxonomic ranks seems to be imminent. This process has already started with the proposal of a phylum rank for-RNA viruses, for which monophyly is unequivocally supported by the present analysis (Figures 1 and 6). This phylum could consist of two subphyla with multiple classes and orders. At least 4 additional phyla of RNA viruses can be confidently predicted to emerge, including, respectively, the dsRNA viruses of Branch 4, and +RNA viruses of Branches 1, 2, and 3. Each of these phyla will undoubtedly have a rich internal structure. In addition, some of the current families do not seem to be compatible with the expansive RdRp trees present here and in a previous analysis (14). For instance, the families *Coronaviridae, Togaviridae* and *Rhabdoviridae* are likely to be split into two families each.

While the present study was in preparation, a major attempt on a comprehensive virus classification has been published (121). This work analyzed a dendrogram that was produced from distance matrices between viruses derived from sequence similarity scores combined with measures of gene composition similarity. Unlike our present analysis, this approach pre-supposes monophyly of each of the Baltimore classes. Furthermore, given that their analysis is based on measures of similarity rather than on phylogenetic analysis proper, this approach is best regarded as producing a phenetic classification of viruses rather that an evolutionary reconstruction as such. Some of the groups delineated by this method, particularly, among +RNA viruses, are closely similar to those reported here. Others, however, are widely different: for instance, the order *Mononegavirales* does not come across as monophyletic in their dendrograms. We did not attempt a complete comparison; such an exercise could be useful in the future, for better understanding the routes of RNA virus evolution.

## CONCLUDING REMARKS

Through metaviromics, many aspects of the global RNA virome evolution can be clarified. Certainly, reconstruction of the deepest events in this evolutionary history is bound to remain tentative, especially, because the RdRp is the only universal gene of the RNA viruses, and hence, the only one that can serve as the template for evolutionary reconstructions. At the depth of divergence characteristic of RdRps, the relationship between the major branches in the tree cannot be established with confidence. Nevertheless, monophyly of several expansive groups, in particular the 5 main branches in the RdRp tree, is strongly supported. Because of the stability of these branches, biologically plausible scenarios of evolution emerge under which dsRNA viruses evolved from different groups of +RNA viruses, whereas -RNA viruses evolved from dsRNA viruses.

Evolutionary reconstructions suggest that the last common ancestors of each major lineage of eukaryotic +RNA viruses were simple viruses that encoded only the RdRp and a CP, most likely, of the SJR fold. Subsequent evolution involved independent capture of distinct helicases which apparently facilitate replication of larger, more complex +RNA genomes. The helicase-assisted replication of +RNA genomes created the opportunities for parallel acquisition of additional genes encoding proteins involved in polyprotein processing and virus genome expression, such as proteases and capping enzymes, respectively, and proteins involved in virus-host interactions, such as MPs or RNAi suppressors of plant viruses. In addition to these processes of vertical evolution of RNA viruses, phylogenomic analysis reveals multiple cases of gene module exchange among diverse viruses and pervasive HVT, often between distantly related hosts, such as animals and plants. Together, these processes have shaped a complex network of evolutionary relationships among RNA viruses.

The much anticipated comprehensive exploration of the RNA viromes of prokaryotes and unicellular eukaryotes, such as free-living excavates, chromalveolates, rhizaria, amoebozoa and choanoflagellates, as well as deeply rooted metazoa, will undoubtedly help in developing better supported evolutionary scenarios for each of the 5 major branches of the RNA virus tree. Nevertheless, it is already clear that the current taxonomy of RNA viruses is due for a complete overhaul.

## MATERIALS AND METHODS

### Phylogeny of RNA-dependent RNA polymerases

Protein sequences belonging to RNA viruses, excluding retroviruses, and unclassified viruses were downloaded from the NCBI GenBank database in April 2017 (122). Initial screening for RdRp domains was performed using PSI-BLAST (123)(e-value of 0.01, effective database size of 2×10^8^) with position-specific scoring matrices (PSSMs) produced from the available RdRp alignments. The sources included group-specific alignments for +RNA viruses and dsRNA viruses (12, 31) and the 4 PFAM alignments for the -RNA viruses (pfam00602, pfam00946, pfam04196, and pfam06317) from the NCBI conserved domain database (CDD) (124). Additionally, a set of RTs from group-II introns and non-long terminal repeat (LTR) retrotransposons was extracted from GenBank as an outgroup.

Extracted RdRp footprints were filtered for the fraction of unresolved amino acids (at most 10%) and clustered using UCLUST (125) with a similarity threshold of 0.9. One representative from each cluster was selected for further analysis. The resulting set contained 4,640 virus RdRps. This set went through several rounds of semi-manual curation whereby sequences were clustered using UCLUST, aligned using MUSCLE (126), and cross-searched against each other and their parent sequences (often, complete viral polyproteins) using PSI-BLAST and HHSEARCH (127). Upon the results of these searches, the boundaries of the RdRp domain were expanded or trimmed to improve their compatibility with each other.

The RdRp and RT sequences were subjected to an iterative clustering and aligned procedure, organized as follows: Initially, sequences were clustered using UCLUST with a similarity threshold of 0.5; clustered sequences were aligned using MUSCLE, and singletons were converted to pseudo-alignments consisting of just one sequence. Sites containing more than 67% of gaps were temporarily removed from alignments and pairwise similarity scores were obtained for clusters using HHSEARCH. Scores for a pair of clusters were converted to distances (the *d*_A,B_ = −log(*s*_A,B_/min(*s*_A,A_, *S*_B,B_) formula, in which *d*_A,B_ is the distance between cluster A and B and *S*_A,B_ is the HHSEARCH score for the comparison of these clusters, was used to convert scores s to distances *d*). The matrix of pairwise distances was used to make an unweighted pair group method with arithmetic (UPGMA) mean (128). Shallow tips of the tree were used as the guide tree for a progressive pairwise alignment of the clusters at the tree leaves using HHALIGN (127), resulting in larger clusters. This procedure was reiterative, ultimately resulting in the single alignment of the whole set of 4,640 virus RdRp sequences and 1,028 RT sequences.

During the clustering procedure, 50 virus RdRp clusters, consisting of 1 to 545 sequences, were defined. These clusters represent either well-established groups of related viruses (roughly comparable to the ICTV family rank) or, in case of poorly characterized and unclassified viruses, groups of well-aligned RdRps that are clearly distinct from others. In all cases, uncertainties were treated conservatively, i.e., in favor of placing sequences with questionable relatedness into separate clusters. Additionally, RT sequences were placed into two clusters consisting of group-II intron RTs and non-LTR retrotransposon RTs.

For each cluster consisting of more than two sequences, an approximate maximum likelihood (ML) phylogenetic tree was constructed from the cluster-specific alignment using the FastTree program (129) (Whelan and Goldman [WAG] evolutionary model with gamma-distributed site rates) with sites, containing more than 50% of gaps removed from the alignment. Trees were rooted using a variant of the mid-point rooting procedure such that the difference between the (weighted) average root-to-tip distances in the subtrees on the opposite sides of the root is minimized.

To resolve the structure of the global relationships, up to five representatives from each cluster were selected using the within-cluster trees to ensure the diversity of the selected sequences. This procedure resulted in a set of 228 virus RdRps and 10 RTs. The alignments of the selected sequences were extracted from the master alignment and filtered for sites containing more than 50% of gaps. A ML phylogenetic tree was reconstructed for the resulting alignment using PhyML (48) (Le Gascuel [LG] evolutionary model with gamma-distributed site rates and empirical amino acid frequencies; aBayes support values). Another form of branch support, bootstrap support by transfer (BOOSTER) phylogenetic bootstrap (130), was also used to assess the reliability of the major tree divisions. Alternatively, the same RdRp alignment was used as the input for ML phylogenetic analysis using RAxML (LG evolutionary model with gamma-distributed site rates and empirical amino acid frequencies).

RdRps in the global tree were divided into 5 major branches (supergroups). Up to 15 representatives from each cluster were selected to form supergroup-level alignments. Respective trees were reconstructed from these alignments using the same procedure (PhyML tree with LG evolutionary model with gamma-distributed site rates and empirical amino acid frequencies).

The overall tree was assembled manually by first replacing the supergroup representatives in the global tree with the supergroup trees, and then, replacing the cluster representatives with the cluster trees. The lower-level trees were rooted according to the arrangement of the representatives in the upper-level tree.

### Identification of protein domains

Protein domains were identified using a representative set of RNA virus genomes, including representative members of all ICTV-approved virus families and unclassified virus groups. This set was annotated manually using sensitive profile-profile comparisons with the HHsuite package (127), and hmm profiles for annotated proteins or their domains were generated by running one iteration of HHblits against the latest (October 2017) uniclust30 database (131). Each annotated profile was assigned to a functional category (e.g., “capsid protein_jelly-roll”, “chymotrypsin-like protease”). These profiles were used to annotate the genomes of all viruses included in the RdRp phylogenetic analysis and for which complete (or near-complete) genome sequences were available. Profiles for the latter proteins were generated by running one iteration of Jackhmmer (132) against UniRef50 database. Protein regions that did not have significant hits were extracted and clustered with cluster analysis of sequences (CLANS) (133), and groups containing at least three members were identified, annotated (if possible), and added to the manually annotated profile database. The last step of the domain identification procedure was then repeated using the updated RNA virus profile database. Highly similar viruses (having identical domain organization and >94% identical RdRps) were removed from the dataset. In addition, only one representative genome was retained for some members of over-represented species (e.g., *Hepacivirus* C). Many genomes encoded readthrough proteins (e.g., alphaviruses, tombusviruses, nidoviruses), resulting in domains from the shorter protein contained within the longer readthrough protein. Such redundancies were removed by filtering out the shorter of the two proteins sharing >80% identity. The final set of annotated genomes included 2,839 viruses.

### Reconstruction of the history of gene gain and loss

The protein domains identified in the 2,839 virus genomes were mapped onto the composite tree of virus RdRps. A set of 500 representatives was chosen among the (mostly complete) genomes. The domain complement data and the respective tree were analyzed using the GLOOME program (51). Domain gains and losses were inferred at each tree branch using the difference of posterior probabilities for the domain occurrence in the nodes at the proximal and the distal ends of the branch (difference >0.5 implies gain; difference <-0.5 implies loss).

### Clustering analysis

The SJR CP network was generated by performing all-against-all comparison of the SJR CP profiles. To generate hmm profiles, SJR CP sequences were extracted from the total proteome of RNA viruses and two iterations of HHblits were performed against the uniprot20_2016_02 database. The resultant profiles were compared to each other with HHsearch (127). Their similarity P-values were extracted from the result files and used as an input for CLANS program (133). Clusters were identified using a network-based algorithm implemented in CLANS. Resulting clusters were manually inspected and refined.

### Analysis of bipartite gene-genome networks

A bipartite network was built to study the patterns of gene sharing among viral genomes (44, 106). After removing genomes with less than 2 domains and domains that appear in less than 3 genomes, the network consisted of 2,829 nodes, of which 2,515 correspond to genomes and 314 to domains. Genome and domain nodes were connected by links whenever a domain is present in a genome. A consensus community detection approach was used to identify the modules of the network (134, 135). First, we ran 500 replicas with Infomap (136) (bipartite setting, using domains as factors) and built a similarity matrix by assigning to each pair of nodes a similarity value equal to the fraction of replicas in which both nodes were placed in the same module. Then, hierarchical clustering was performed on the similarity matrix, setting the number of clusters equal to the median number of modules obtained in the 500 replicas. The order statistics local optimization method (Oslom) software (137) was subsequently used to filter significant modules with a p-value threshold equal to 0.05. To detect higher-order (super)modules, we first identified connector domains as those present in at least 2 modules with prevalence greater than 0.65. The 2nd-order network composed of 54 modules and 34 connector domains was searched for 2nd-order modules with Infomap (500 replicas, bipartite setting with connector domains as factors). Due to the small size of the 2nd-order network, consensus community detection did not qualitatively improve the results of the search, and therefore we took the replica with the best Infomap score and skipped hierarchical clustering for this and subsequent steps of the higher-order module search. After assessing statistical significance with Oslom, 4 2nd-order modules were recovered, encompassing 12 of the original modules associated with closely related virus families. A 3rd-order network was built by pooling these 4 2nd-order modules withthe 46 modules that remained unmerged. The 3rd-order network was analyzed in the same manner to obtain the 9 supermodules. None of these supermodules was assessed as significant by Oslom.

### Quantification of virus host range diversity

Known hosts or, in case of viruses, isolated from holobionts, virus sources were identified for 3,456 viruses. The host taxonomy (to the phylum level) was mapped to the leaves of the combined tree. The tree was ultrametrized and the weights were computed for all leaves as described in (138). Each internal node of the tree, therefore, defines a set of leaves with assigned hosts; the distribution of (weighted) relative frequencies of hosts can be characterized by its Shannon entropy. Each leaf has one host defined, so the entropy of the host range distribution is 0, whereas at the root the host diversity is maximal. The entropy generally declines along the tree because viruses that belong to relatively shallow branches tend to share their host ranges. The node depth vs node host range entropy data was averaged for each major branch separately using a Gaussian kernel with the bandwidth equal to 10% of the total tree depth.

## ACKNOWLEDGMENTS

The content of this publication does not necessarily reflect the views or policies of the US Department of the Army, the US Department of Defense, or the US Department of Health and Human Services or of the institutions and companies affiliated with the authors. VVD is grateful to Barbara and Simon Gvakharia for their support and inspiration. YIW and EVK are supported through the intramural program of the National Institutes of Health of the USA. DK was partly supported by a Short Term Fellowship from the Federation of European Biochemical Societies (FEBS). MK was supported by l’Agence Nationale de la Recherche (ANR, France) project ENVIRA. This work was funded in part through Battelle Memorial Institute’s prime contract with the US National Institute of Allergy and Infectious Diseases (NIAID) under Contract No. HHSN272200700016I (JHK). We thank Jiro Wada (IRF-Frederick) for help with figure creation.

## SUPPLEMENTAL MATERIAL

Supplemental Figure S1. Iterative clustering-alignment-phylogeny procedure.

Supplemental Figure S2. (A) Gain and loss of capsid (nucleocapsid) proteins and movement proteins. (B) Gain and loss of key enzymes. Each branch is shown by a triangle which represents collapsed sequences of the respective set of RdRps. The 5 main branches discussed in the text are indicated. The genes (domains) are denoted by distinct shapes shown at the bottom of each panel. Different colors show distinct families of proteins with similar functions (helicases, proteases, capping enzymes) and different hues of the same color show distinct subfamilies of the same family that are likely to have been acauired independently. The symbols at internal nodes denoted the inferred point of origin (gain) of the respective gene. Empty shapes show loss of the respective genes. Shapes placed inside triangles indicate the presence of the respective gene in a subset of the viruses in the respective branch.

Supplemental Figure S3. Sequence similarity networks of SJR-CPs. Protein sequences were clustered by the pairwise similarity of their hmm profiles using CLANS. Different groups of SJR-CPs are shown as clouds of differentially colored circles, with the corresponding subgroups labeled as indicated in the figure. Edges connect sequences with CLANS P-value < 1e-10.

Supplemental Data set S1. List of viruses.

Supplemental Data set S2. Phylogenetic tree for the global set of RdRps; Newick format.

Supplemental Data set S3. Phylogenetic trees for the global set of RdRp representatives; Newick format. A. Reconstructed using PhyML. B. Reconstructed using RAxML.

Supplemental Data set S4. Phylogenetic trees for RdRp representatives; Newick format.

A. Branch 1 representatives.

B. Branch 2 representatives.

C. Branch 3 representatives.

D. Branch 4 representatives.

E. Branch 5 representatives

Supplemental Data set S5. Proteins domains encoded by the virus genomes.

Additional data can be found at the following directory: ftp://ftp.ncbi.nlm.nih.gov/pub/wolf/suppl/rnavir18

## REFERENCES

1 Bernhardt HS. 2012. The RNA world hypothesis: the worst theory of the early evolution of life (except for all the others)(a). Biol Direct 7: 23.

2 Gilbert W. 1986. Origin of Life the RNA World. Nature 319: 618–618.

3 Nelson JW, Breaker RR. 2017. The lost language of the RNA World. Sci Signal 10: eaam8812.

4 Koonin EV, Senkevich TG, Dolja VV. 2006. The ancient Virus World and evolution of cells. Biol Direct 1: 29.

5 Baltimore D. 1971. Expression of animal virus genomes. Bacteriol Rev 35: 235–241.

6 Baltimore D. 1971. Viral genetic systems. Trans N Y Acad Sci 33: 327–332.

7 Koonin EV, Dolja VV. 1993. Evolution and taxonomy of positive-strand RNA viruses: implications of comparative analysis of amino acid sequences. Crit Rev Biochem Mol Biol 28: 375–430.

8 Koonin EV, Dolja VV. 2013. A virocentric perspective on the evolution of life. Curr Opin Virol 3: 546–557.

9 Ahlquist P. 2006. Parallels among positive-strand RNA viruses, reverse-transcribing viruses and double-stranded RNA viruses. Nat Rev Microbiol 4: 371–382.

10 Reguera J, Gerlach P, Cusack S. 2016. Towards a structural understanding of RNA synthesis by negative strand RNA viral polymerases. Curr Opin Struct Biol 36: 75–84.

11 Dolja VV, Koonin EV. 2018. Metagenomics reshapes the concepts of RNA virus evolution by revealing extensive horizontal virus transfer. Virus Res 244: 36–52.

12 Bolduc B, Shaughnessy DP, Wolf YI, Koonin EV, Roberto FF, Young M. 2012. Identification of novel positive-strand RNA viruses by metagenomic analysis of archaea-dominated Yellowstone hot springs. J Virol 86: 5562–5573.

13 Krishnamurthy SR, Janowski AB, Zhao G, Barouch D, Wang D. 2016. Hyperexpansion of RNA bacteriophage diversity. PLoS Biol 14: e1002409.

14 Shi M, Lin X-D, Tian J-H, Chen L-J, Chen X, Li C-X, Qin X-C, Li J, Cao J-P, Eden J-S, Buchmann J, Wang W, Xu J, Holmes EC, Zhang Y-Z. 2016. Redefining the invertebrate RNA virosphere. Nature 540: 539–543.

15 Koonin EV, Dolja VV, Krupovic M. 2015. Origins and evolution of viruses of eukaryotes: the ultimate modularity. Virology 479–480: 2–25.

16 Dolja VV, Koonin EV. 2011. Common origins and host-dependent diversity of plant and animal viromes. Curr Opin Virol 1: 322–331.

17 Shi M, Zhang Y-Z, Holmes EC. 2018. Meta-transcriptomics and The Evolutionary Biology of RNA Viruses. Virus Res 243: 83–90.

18 Koonin EV, Dolja VV. 2014. Virus world as an evolutionary network of viruses and capsidless selfish elements. Microbiol Mol Biol Rev 78: 278–303.

19 Krupovic M, Koonin EV. 2017. Multiple origins of viral capsid proteins from cellular ancestors. Proc Natl Acad Sci U S A 114: E2401–E2410.

20 Dolja VV, Boyko VP, Agranovsky AA, Koonin EV. 1991. Phylogeny of capsid proteins of rod-shaped and filamentous RNA plant viruses: two families with distinct patterns of sequence and probably structure conservation. Virology 184: 79–86.

21 Koonin EV. 1991. The phylogeny of RNA-dependent RNA polymerases of positive-strand RNA viruses. J Gen Virol 72 (Pt 9): 2197–2206.

22 Xiong Y, Eickbush TH. 1990. Origin and evolution of retroelements based upon their reverse transcriptase sequences. EMBO J 9: 3353–3362.

23 Poch O, Sauvaget I, Delarue M, Tordo N. 1989. Identification of four conserved motifs among the RNA-dependent polymerase encoding elements. EMBO J 8: 3867–3874.

24 Ng KK-S, Arnold JJ, Cameron CE. 2008. Structure-function relationships among RNA-dependent RNA polymerases. Curr Top Microbiol Immunol 320: 137–156.

25 te Velthuis AJW. 2014. Common and unique features of viral RNA-dependent polymerases. Cell Mol Life Sci 71: 4403–4420.

26 Eickbush TH, Jamburuthugoda VK. 2008. The diversity of retrotransposons and the properties of their reverse transcriptases. Virus Res 134: 221–234.

27 Gladyshev EA, Arkhipova IR. 2011. A widespread class of reverse transcriptase-related cellular genes. Proc Natl Acad Sci U S A 108: 20311–20316.

28 Lambowitz AM, Belfort M. 2015. Mobile bacterial group II introns at the crux of eukaryotic evolution. Microbiol Spectr 3: MDNA3–0050–2014.

29 Novikova O, Belfort M. 2017. Mobile group II introns as ancestral eukaryotic elements. Trends Genet 33: 773–783.

30 Krupovic M, Blomberg J, Coffin JM, Dasgupta I, Fan H, Geering AD, Gifford R, Harrach B, Hull R, Johnson W, Kreuze JF, Lindemann D, Llorens C, Lockhart B, Mayer J, Muller E, Olszewski NE, Pappu HR, Pooggin MM, Richert-Poggeler KR, Sabanadzovic S, Sanfacon H, Schoelz JE, Seal S, Stavolone L, Stoye JP, Teycheney P-Y, Tristem M, Koonin EV, Kuhn JH. 2018. Ortervirales: new virus order unifying five families of reverse-transcribing viruses. J Virol 92: e00515–00518.

31 Koonin EV, Wolf YI, Nagasaki K, Dolja VV. 2008. The Big Bang of picorna-like virus evolution antedates the radiation of eukaryotic supergroups. Nat Rev Microbiol 6: 925–939.

32 Koonin EV. 1992. Evolution of double-stranded RNA viruses: a case for polyphyletic origin from different groups of positive-stranded RNA viruses. Semin Virol 3: 327–339.

33 El Omari K, Sutton G, Ravantti JJ, Zhang H, Walter TS, Grimes JM, Bamford DH, Stuart DI, Mancini EJ. 2013. Plate tectonics of virus shell assembly and reorganization in phage Φ8, a distant relative of mammalian reoviruses. Structure 21: 1384–1395.

34 Poranen MM, Bamford DH. 2012. Assembly of large icosahedral double-stranded RNA viruses. Adv Exp Med Biol 726: 379–402.

35 Pflug A, Guilligay D, Reich S, Cusack S. 2014. Structure of influenza A polymerase bound to the viral RNA promoter. Nature 516: 355–360.

36 Goldbach R, Wellink J. 1988. Evolution of plus-strand RNA viruses. Intervirology 29: 260–267.

37 Greninger AL. 2018. A decade of RNA virus metagenomics is (not) enough. Virus Res 244: 218–229.

38 Shi M, Lin X-D, Vasilakis N, Tian J-H, Li C-X, Chen L-J, Eastwood G, Diao X-N, Chen M-H, Chen X, Qin X-C, Widen SG, Wood TG, Tesh RB, Xu J, Holmes EC, Zhang Y-Z. 2016. Divergent viruses discovered in arthropods and vertebrates revise the evolutionary history of the *Flaviviridae* and related viruses. J Virol 90: 659–669.

39 Li C-X, Shi M, Tian J-H, Lin X-D, Kang Y-J, Chen L-J, Qin X-C, Xu J, Holmes EC, Zhang Y-Z. 2015. Unprecedented genomic diversity of RNA viruses in arthropods reveals the ancestry of negative-sense RNA viruses. Elife 4: e05378.

40 Webster CL, Longdon B, Lewis SH, Obbard DJ. 2016. Twenty-five new viruses associated with the Drosophilidae (Diptera). Evol Bioinform Online 12: 13–25.

41 Fauver JR, Grubaugh ND, Krajacich BJ, Weger-Lucarelli J, Lakin SM, Fakoli LS, 3rd, Bolay FK, Diclaro JW, 2nd, Dabire KR, Foy BD, Brackney DE, Ebel GD, Stenglein MD. 2016. West African *Anopheles gambiae* mosquitoes harbor a taxonomically diverse virome including new insect-specific flaviviruses, mononegaviruses, and totiviruses. Virology 498: 288–299.

42 Marzano S-YL, Nelson BD, Ajayi-Oyetunde O, Bradley CA, Hughes TJ, Hartman GL, Eastburn DM, Domier LL. 2016. Identification of diverse mycoviruses through metatranscriptomics characterization of the viromes of five major fungal plant pathogens. J Virol 90: 6846–6863.

43 Deakin G, Dobbs E, Bennett JM, Jones IM, Grogan HM, Burton KS. 2017. Multiple viral infections in *Agaricus bisporus*-Characterisation of 18 unique RNA viruses and 8 ORFans identified by deep sequencing. Sci Rep 7: 2469.

44 Iranzo J, Krupovic M, Koonin EV. 2016. The double-stranded DNA virosphere as a modular hierarchical network of gene sharing. MBio 7: e00978–00916.

45 Iranzo J, Krupovic M, Koonin EV. 2017. A network perspective on the virus world. Commun Integr Biol 10: e1296614.

46 Gorbalenya AE, Pringle FM, Zeddam J-L, Luke BT, Cameron CE, Kalmakoff J, Hanzlik TN, Gordon KHJ, Ward VK. 2002. The palm subdomain-based active site is internally permuted in viral RNA-dependent RNA polymerases of an ancient lineage. J Mol Biol 324: 47–62.

47 King AMQ, Lefkowitz EJ, Mushegian AR, Adams MJ, Dutilh BE, Gorbalenya AE, Harrach B, Harrison RL, Junglen S, Knowles NJ, Kropinski AM, Krupovic M, Kuhn JH, Nibert ML, Rubino L, Sabanadzovic S, Sanfaçon H, Siddell SG, Simmonds P, Varsani A, Zerbini FM, Davison AJ. 2018. Changes to taxonomy and the International Code of Virus Classification and Nomenclature ratified by the International Committee on Taxonomy of Viruses (2018). Arch Virol 163: 2601–2631.

48 Guindon S, Dufayard J-F, Lefort V, Anisimova M, Hordijk W, Gascuel O. 2010. New algorithms and methods to estimate maximum-likelihood phylogenies: assessing the performance of PhyML 3.0. Syst Biol 59: 307–321.

49 Akopyants NS, Lye L-F, Dobson DE, Lukeš J, Beverley SM. 2016. A novel bunyavirus-like virus of trypanosomatid protist parasites. Genome Announc 4: e00715–00716.

50 Grybchuk D, Akopyants NS, Kostygov AY, Konovalovas A, Lye L-F, Dobson DE, Zangger H, Fasel N, Butenko A, Frolov AO, Votýpka J, d’Avila-Levy CM, Kulich P, Moravcová J, Plevka P, Rogozin IB, Serva S, Lukeš J, Beverley SM, Yurchenko V. 2018. Viral discovery and diversity in trypanosomatid protozoa with a focus on relatives of the human parasite Leishmania. Proc Natl Acad Sci U S A 115: E506–E515.

51 Cohen O, Ashkenazy H, Belinky F, Huchon D, Pupko T. 2010. GLOOME: gain loss mapping engine. Bioinformatics 26: 2914–2915.

52 Golmohammadi R, Valegård K, Fridborg K, Liljas L. 1993. The refined structure of bacteriophage MS2 at 2.8 Å resolution. J Mol Biol 234: 620–639.

53 Hillman BI, Cai G. 2013. The family Narnaviridae: simplest of RNA viruses. Adv Virus Res 86: 149–176.

54 Nibert ML, Vong M, Fugate KK, Debat HJ. 2018. Evidence for contemporary plant mitoviruses. Virology 518: 14–24.

55 Rastgou M, Habibi MK, Izadpanah K, Masenga V, Milne RG, Wolf YI, Koonin EV, Turina M. 2009. Molecular characterization of the plant virus genus *Ourmiavirus* and evidence of inter-kingdom reassortment of viral genome segments as its possible route of origin. J Gen Virol 90: 2525–2535.

56 Le Gall O, Christian P, Fauquet CM, King AMQ, Knowles NJ, Nakashima N, Stanway G, Gorbalenya AE. 2008. Picornavirales, a proposed order of positive-sense single-stranded RNA viruses with a pseudo-T = 3 virion architecture. Arch Virol 153: 715–727.

57 Culley A. 2018. New insight into the RNA aquatic virosphere via viromics. Virus Research 244: 84–89.

58 Ng TFF, Marine R, Wang C, Simmonds P, Kapusinszky B, Bodhidatta L, Oderinde BS, Wommack KE, Delwart E. 2012. High variety of known and new RNA and DNA viruses of diverse origins in untreated sewage. J Virol 86: 12161–12175.

59 Hause BM, Palinski R, Hesse R, Anderson G. 2016. Highly diverse posaviruses in swine faeces are aquatic in origin. J Gen Virol 97: 1362–1367.

60 Jiang B, Monroe SS, Koonin EV, Stine SE, Glass RI. 1993. RNA sequence of astrovirus: distinctive genomic organization and a putative retrovirus-like ribosomal frameshifting signal that directs the viral replicase synthesis. Proc Natl Acad Sci U S A 90: 10539–10543.

61 Saberi A, Gulyaeva AA, Brubacher JL, Newmark PA, Gorbalenya AE. 2018. A planarian nidovirus expands the limits of RNA genome size. bioRxiv doi: 10.1101/299776:299776.

62 Nagasaki K, Shirai Y, Takao Y, Mizumoto H, Nishida K, Tomaru Y. 2005. Comparison of genome sequences of single-stranded RNA viruses infecting the bivalvekilling dinoflagellate Heterocapsa circularisquama. Appl Environ Microbiol 71: 8888–8894.

63 Koonin EV, Choi GH, Nuss DL, Shapira R, Carrington JC. 1991. Evidence for common ancestry of a chestnut blight hypovirulence-associated double-stranded RNA and a group of positive-strand RNA plant viruses. Proc Natl Acad Sci U S A 88: 10647–10651.

64 Dawe AL, Nuss DL. 2013. Hypovirus molecular biology: from Koch’s postulates to host self-recognition genes that restrict virus transmission. Adv Virus Res 86: 109–147.

65 Nibert ML, Ghabrial SA, Maiss E, Lesker T, Vainio EJ, Jiang D, Suzuki N. 2014. Taxonomic reorganization of family *Partitiviridae* and other recent progress in partitivirus research. Virus Res 188: 128–141.

66 Tang J, Ochoa WF, Li H, Havens WM, Nibert ML, Ghabrial SA, Baker TS. 2010. Structure of Fusarium poae virus 1 shows conserved and variable elements of partitivirus capsids and evolutionary relationships to picobirnavirus. J Struct Biol 172: 363–371.

67 Duquerroy S, Da Costa B, Henry C, Vigouroux A, Libersou S, Lepault J, Navaza J, Delmas B, Rey FA. 2009. The picobirnavirus crystal structure provides functional insights into virion assembly and cell entry. EMBO J 28: 1655–1665.

68 Nibert ML, Tang J, Xie J, Collier AM, Ghabrial SA, Baker TS, Tao YJ. 2013. 3D structures of fungal partitiviruses. Adv Virus Res 86: 59–85.

69 Luque D, Gómez-Blanco J, Garriga D, Brilot AF, González JM, Havens WM, Carrascosa JL, Trus BL, Verdaguer N, Ghabrial SA, Castón JR. 2014. Cryo-EM near-atomic structure of a dsRNA fungal virus shows ancient structural motifs preserved in the dsRNA viral lineage. Proc Natl Acad Sci U S A 111: 7641–7646.

70 Koga R, Fukuhara T, Nitta T. 1998. Molecular characterization of a single mitochondria-associated double-stranded RNA in the green alga Bryopsis. Plant Mol Biol 36: 717–724.

71 Koga R, Horiuchi H, Fukuhara T. 2003. Double-stranded RNA replicons associated with chloroplasts of a green alga *Bryopsis cinicola*. Plant Mol Biol 51: 991–999.

72 Koonin EV, Dolja VV. 2006. Evolution of complexity in the viral world: the dawn of a new vision. Virus Res 117: 1–4.

73 Gorbalenya AE, Koonin EV. 1989. Viral proteins containing the purine NTP-binding sequence pattern. Nucleic Acids Res 17: 8413–8440.

74 Snijder EJ, Decroly E, Ziebuhr J. 2016. The nonstructural proteins directing coronavirus RNA synthesis and processing. Adv Virus Res 96: 59–126.

75 Enjuanes L, Zuñiga S, Castaño-Rodriguez C, Gutierrez-Alvarez J, Canton J, Sola I. 2016. Molecular basis of coronavirus virulence and vaccine development. Adv Virus Res 96: 245–286.

76 Sola I, Almazán F, Zúñiga S, Enjuanes L. 2015. Continuous and discontinuous RNA synthesis in coronaviruses. Annu Rev Virol 2: 265–288.

77 Roossinck MJ, Sabanadzovic S, Okada R, Valverde RA. 2011. The remarkable evolutionary history of endornaviruses. J Gen Virol 92: 2674–2678.

78 Li K, Zheng D, Cheng J, Chen T, Fu Y, Jiang D, Xie J. 2016. Characterization of a novel *Sclerotinia sclerotiorum* RNA virus as the prototype of a new proposed family within the order Tymovirales. Virus Res 219: 92–99.

79 Syller J. 2002. Umbraviruses-the unique plant viruses that do not encode a capsid protein. Acta Microbiol Pol 51: 99–113.

80 Janowski AB, Krishnamurthy SR, Lim ES, Zhao G, Brenchley JM, Barouch DH, Thakwalakwa C, Manary MJ, Holtz LR, Wang D. 2017. Statoviruses, a novel taxon of RNA viruses present in the gastrointestinal tracts of diverse mammals. Virology 504: 36–44.

81 Greninger AL, DeRisi JL. 2015. Draft genome sequence of tombunodavirus UC1. Genome Announc 3: e00655–00615.

82 Dokland T, Walsh M, Mackenzie JM, Khromykh AA, Ee K-H, Wang S. 2004. West Nile virus core protein; tetramer structure and ribbon formation. Structure 12: 1157–1163.

83 Ma L, Jones CT, Groesch TD, Kuhn RJ, Post CB. 2004. Solution structure of dengue virus capsid protein reveals another fold. Proc Natl Acad Sci U S A 101: 3414–3419.

84 Kuhn RJ, Rossmann MG. 2005. Structure and assembly of icosahedral enveloped RNA viruses. Adv Virus Res 64: 263–284.

85 Ahola T, Karlin DG. 2015. Sequence analysis reveals a conserved extension in the capping enzyme of the alphavirus supergroup, and a homologous domain in nodaviruses. Biol Direct 10: 16.

86 Koonin EV. 1993. Computer-assisted identification of a putative methyltransferase domain in NS5 protein of flaviviruses and λ2 protein of reovirus. J Gen Virol 74 (Pt 4): 733–740.

87 Liu L, Dong H, Chen H, Zhang J, Ling H, Li Z, Shi P-Y, Li H. 2010. Flavivirus RNA cap methyltransferase: structure, function, and inhibition. Front Biol (Beijing) 5: 286–303.

88 Dolja VV, Kreuze JF, Valkonen JPT. 2006. Comparative and functional genomics of closteroviruses. Virus Res 117: 38–51.

89 Kobayashi K, Atsumi G, Iwadate Y, Tomita R, Chiba K-I, Akasaka S, Nishihara M, Takahashi H, Yamaoka N, Nishiguchi M, Sekine K-T. 2013. Gentian Kobu-sho-associated virus: a tentative, novel double-stranded RNA virus that is relevant to gentian Kobu-sho syndrome. J Gen Plant Pathol 79: 56–63.

90 Teixeira M, Sela N, Ng J, Casteel CL, Peng H-C, Bekal S, Girke T, Ghanim M, Kaloshian I. 2016. A novel virus from *Macrosiphum euphorbiae* with similarities to members of the family Flaviviridae. J Gen Virol 97: 1261–1271.

91 Mata CP, Luque D, Gómez-Blanco J, Rodríguez JM, González JM, Suzuki N, Ghabrial SA, Carrascosa JL, Trus BL, Castón JR. 2017. Acquisition of functions on the outer capsid surface during evolution of double-stranded RNA fungal viruses. PLoS Pathog 13: e1006755.

92 Castón JR, Luque D, Gómez-Blanco J, Ghabrial SA. 2013. Chrysovirus structure: repeated helical core as evidence of gene duplication. Adv Virus Res 86: 87–108.

93 Pan J, Dong L, Lin L, Ochoa WF, Sinkovits RS, Havens WM, Nibert ML, Baker TS, Ghabrial SA, Tao YJ. 2009. Atomic structure reveals the unique capsid organization of a dsRNA virus. Proc Natl Acad Sci U S A 106: 4225–4230.

94 Abrescia NGA, Bamford DH, Grimes JM, Stuart DI. 2012. Structure unifies the viral universe. Annu Rev Biochem 81: 795–822.

95 El Omari K, Meier C, Kainov D, Sutton G, Grimes JM, Poranen MM, Bamford DH, Tuma R, Stuart DI, Mancini EJ. 2013. Tracking in atomic detail the functional specializations in viral RecA helicases that occur during evolution. Nucleic Acids Res 41: 9396–9410.

96 Sutton G, Grimes JM, Stuart DI, Roy P. 2007. Bluetongue virus VP4 is an RNA-capping assembly line. Nat Struct Mol Biol 14: 449–451.

97 Yu X, Jiang J, Sun J, Zhou ZH. 2015. A putative ATPase mediates RNA transcription and capping in a dsRNA virus. Elife 4: e07901.

98 Amarasinghe GK, Ceballos NGA, Banyard AC, Basler CF, Bavari S, Bennett AJ, Blasdell KR, Briese T, Bukreyev A, Caì Y, Calisher CH, Lawson CC, Chandran K, Chapman CA, Chiu CY, Choi K-S, Collins PL, Dietzgen RG, Dolja VV, Dolnik O, Domier LL, Dürrwald R, Dye JM, Easton AJ, Ebihara H, Echevarría JE, Fooks AR, Formenty PBH, Fouchier RAM, Freuling CM, Ghedin E, Goldberg TL, Hewson R, Horie M, Hyndman TH, Jiāng D, Kityo R, Kobinger GP, Kondō H, Koonin EV, Krupovic M, Kurath G, Lamb RA, Lee B, Leroy EM, Maes P, Maisner A, Marston DA, Mor SK, Müller T, et al. 2018. Taxonomy of the order *Mononegavirales*: update 2018. Arch Virol 163: 2283–2294.

99 Kormelink R, Garcia ML, Goodin M, Sasaya T, Haenni A-L. 2011. Negative-strand’ RNA viruses: the plant-infecting counterparts. Virus Res 162: 184–202.

100 Osaki H, Sasaki A, Nomiyama K, Tomioka K. 2016. Multiple virus infection in a single strain of *Fusarium poae* shown by deep sequencing. Virus Genes 52: 835–847.

101 Bacharach E, Mishra N, Briese T, Zody MC, Kembou Tsofack JE, Zamostiano R, Berkowitz A, Ng J, Nitido A, Corvelo A, Toussaint NC, Abel Nielsen SC, Hornig M, Del Pozo J, Bloom T, Ferguson H, Eldar A, Lipkin WI. 2016. Characterization of a novel orthomyxo-like virus causing mass die-offs of tilapia. MBio 7: e00431–00416.

102 Maes P, Alkhovsky SV, Bío Y, Beer M, Birkhead M, Briese T, Buchmeier MJ, Calisher CH, Charrel RN, Choi IR, Clegg CS, Torre JCdl, Delwart E, DeRisi JL, Bello PLD, Serio FD, Digiaro M, Dolja VV, Drosten C, Druciarek TZ, Du J, Ebihara H, Elbeaino T, Gergerich RC, Gillis AN, Gonzalez J-PJ, Haenni A-L, Hepojoki J, Hetzel U, Hồ T, Hóng N, Jain RK, Vuren PJv, Jin Q, Jonson MG, Junglen S, Keller KE, Kemp A, Kipar A, Kondov NO, Koonin EV, Kormelink R, Korzyukov Y, Krupovic M, Lambert AJ, Laney AG, LeBreton M, Lukashevic IS, Marklewitz M, Markotter W, et al. 2018. Taxonomy of the family Arenaviridae and the order Bunyavirales: update 2018. Arch Virol 163: 2295–2310.

103 Mielke-Ehret N, Mühlbach H-P. 2012. Emaravirus: a novel genus of multipartite, negative strand RNA plant viruses. Viruses 4: 1515–1536.

104 Krupovic M. 2013. Networks of evolutionary interactions underlying the polyphyletic origin of ssDNA viruses. Curr Opin Virol 3: 578–586.

105 Kazlauskas D, Varsani A, Krupovic M. 2018. Pervasive chimerism in the replication-associated proteins of uncultured single-stranded DNA viruses. Viruses 10: 187.

106 Iranzo J, Koonin EV, Prangishvili D, Krupovic M. 2016. Bipartite network analysis of the archaeal virosphere: evolutionary connections between viruses and capsidless mobile elements. J Virol 90: 11043–11055.

107 Krishnamurthy SR, Wang D. 2018. Extensive conservation of prokaryotic ribosomal binding sites in known and novel picobirnaviruses. Virology 516: 108–114.

108 Boros Á, Polgár B, Pankovics P, Fenyvesi H, Engelmann P, Phan TG, Delwart E, Reuter G. 2018. Multiple divergent picobirnaviruses with functional prokaryotic Shine-Dalgarno ribosome binding sites present in cloacal sample of a diarrheic chicken. Virology 525: 62–72.

109 Jinek M, Doudna JA. 2009. A three-dimensional view of the molecular machinery of RNA interference. Nature 457: 405–412.

110 Wu Q, Wang X, Ding S-W. 2010. Viral suppressors of RNA-based viral immunity: host targets. Cell Host Microbe 8: 12–15.

111 Krupovic M, Dolja VV, Koonin EV. 2015. Plant viruses of the *Amalgaviridae* family evolved via recombination between viruses with double-stranded and negative-strand RNA genomes. Biol Direct 10: 12.

112 Martin RR, Zhou J, Tzanetakis IE. 2011. Blueberry latent virus: an amalgam of the *Partitiviridae and Totiviridae*. Virus Res 155: 175–180.

113 Sabanadzovic S, Valverde RA, Brown JK, Martin RR, Tzanetakis IE. 2009. Southern tomato virus: the link between the families *Totiviridae and Partitiviridae*. Virus Res 140: 130–137.

114 Sabanadzovic S, Abou Ghanem-Sabanadzovic N, Valverde RA. 2010. A novel monopartite dsRNA virus from rhododendron. Arch Virol 155: 1859–1863.

115 Mushegian AR, Koonin EV. 1993. Cell-to-cell movement of plant viruses. Insights from amino acid sequence comparisons of movement proteins and from analogies with cellular transport systems. Arch Virol 133: 239–257.

116 Ghabrial SA, Castón JR, Jiang D, Nibert ML, Suzuki N. 2015. 50-plus years of fungal viruses. Virology 479–480: 356–368.

117 Hillman BI, Annisa A, Suzuki N. 2018. Viruses of plant-interacting fungi. Adv Virus Res 100: 99–116.

118 Kotta-Loizou I, Coutts RHA. 2017. Studies on the virome of the entomopathogenic fungus *Beauveria bassiana* reveal novel dsRNA elements and mild hypervirulence. PLoS Pathog 13: e1006183.

119 Coyle MC, Elya CN, Bronski M, Eisen MB. 2018. Entomophthovirus: an insect-derived iflavirus that infects a behavior manipulating fungal pathogen of dipterans. bioRxiv doi: 10.1101/371526:371526.

120 Waldron FM, Stone GN, Obbard DJ. 2018. Metagenomic sequencing suggests a diversity of RNA interference-like responses to viruses across multicellular eukaryotes. PLoS Genet 14: e1007533.

121 Aiewsakun P, Simmonds P. 2018. The genomic underpinnings of eukaryotic virus taxonomy: creating a sequence-based framework for family-level virus classification. Microbiome 6: 38.

122 Benson DA, Cavanaugh M, Clark K, Karsch-Mizrachi I, Ostell J, Pruitt KD, Sayers EW. 2018. GenBank. Nucleic Acids Res 46: D41–D47.

123 Altschul SF, Madden TL, Schäffer AA, Zhang J, Zhang Z, Miller W, Lipman DJ. 1997. Gapped BLAST and PSI-BLAST: a new generation of protein database search programs. Nucleic Acids Res 25: 3389–3402.

124 Marchler-Bauer A, Derbyshire MK, Gonzales NR, Lu S, Chitsaz F, Geer LY, Geer RC, He J, Gwadz M, Hurwitz DI, Lanczycki CJ, Lu F, Marchler GH, Song JS, Thanki N, Wang Z, Yamashita RA, Zhang D, Zheng C, Bryant SH. 2015. CDD: NCBI’s conserved domain database. Nucleic Acids Res 43: D222–226.

125 Edgar RC. 2010. Search and clustering orders of magnitude faster than BLAST. Bioinformatics 26: 2460–2461.

126 Edgar RC. 2004. MUSCLE: a multiple sequence alignment method with reduced time and space complexity. BMC Bioinformatics 5: 113.

127 Soding J. 2005. Protein homology detection by HMM-HMM comparison. Bioinformatics 21: 951–960.

128 Sokal RB, Michener CD. 1958. A statistical method for evaluating systematic relationships. Univ Kansas Sci Bull XXXVIII: 1409–1438.

129 Price MN, Dehal PS, Arkin AP. 2010. FastTree 2-approximately maximum-likelihood trees for large alignments. PLoS One 5: e9490.

130 Lemoine F, Domelevo Entfellner J-B, Wilkinson E, Correia D, Dávila Felipe M, De Oliveira T, Gascuel O. 2018. Renewing Felsenstein’s phylogenetic bootstrap in the era of big data. Nature 556: 452–456.

131 Mirdita M, von den Driesch L, Galiez C, Martin MJ, Söding J, Steinegger M. 2017. Uniclust databases of clustered and deeply annotated protein sequences and alignments. Nucleic Acids Res 45: D170–D176.

132 Eddy SR. 2011. Accelerated profile HMM searches. PLoS Comput Biol 7: e1002195.

133 Frickey T, Lupas A. 2004. CLANS: a Java application for visualizing protein families based on pairwise similarity. Bioinformatics 20: 3702–3704.

134 Lancichinetti A, Fortunato S. 2012. Consensus clustering in complex networks. Sci Rep 2: 336.

135 Fortunato S, Hric D. 2016. Community detection in networks: A user guide. Physics Rep Rev Sect Physics Lett 659: 1–44.

136 Rosvall M, Bergstrom CT. 2008. Maps of random walks on complex networks reveal community structure. Proc Natl Acad Sci U S A 105: 1118–1123.

137 Lancichinetti A, Radicchi F, Ramasco JJ, Fortunato S. 2011. Finding statistically significant communities in networks. PLoS One 6: e18961.

138 Petitjean C, Makarova KS, Wolf YI, Koonin EV. 2017. Extreme deviations from expected evolutionary rates in archaeal protein families. Genome Biol Evol 9: 2791–2811.

